# A plant pathogen utilizes effector proteins for microbiome manipulation

**DOI:** 10.1101/2020.01.30.926725

**Authors:** Nick C. Snelders, Hanna Rovenich, Gabriella C. Petti, Mercedes Rocafort, Julia A. Vorholt, Jeroen R. Mesters, Michael F. Seidl, Reindert Nijland, Bart P.H.J. Thomma

## Abstract

During colonization of their hosts, pathogens secrete effector proteins to promote disease development through various mechanisms. Increasing evidence shows that the host microbiome plays a crucial role in health, and that hosts actively shape their microbiomes to suppress disease. We hypothesized that pathogens evolved to manipulate host microbiomes to their advantage in turn. Here, we show that the fungal plant pathogen *Verticillium dahliae* utilizes effector proteins for niche colonization through selective manipulation of host microbiomes by suppressing microbes with antagonistic activities. Moreover, we show that effector proteins are similarly exploited for microbiome manipulation in the soil environment, where the fungus resides in absence of a host. In conclusion, we demonstrate that pathogens utilize effector proteins to modulate microbiome compositions and propose that their effector catalogs represent an untapped resource for novel antibiotics.

## Introduction

To establish disease, pathogenic microbes secrete a wide diversity of effector proteins that facilitate host colonization through a multitude of mechanisms^1^. Typically, pathogen effectors are defined as small cysteine-rich proteins that are secreted upon colonization to manipulate host physiology or to deregulate host immune responses^2^. Consequently, effector proteins are predominantly studied in binary host-microbe interactions, while largely ignoring the biotic context in which these interactions take place. Higher organisms, including plants, associate with a plethora of microbes that collectively form their microbiome, which represents a key determinant for their health^3–7^. The most extensive microbial colonization of plants occurs at roots, where plants define rhizosphere microbiome compositions through secretion of exudates^8,9^ and specifically attract beneficial microbes to suppress pathogen invasion^10–12^. Thus, we hypothesized that plant pathogens evolved mechanisms to counteract this recruitment and modulate host microbiomes for successful infection, possibly through effector proteins^1,13^.

*Verticillium dahliae* is a soil-borne fungus that causes vascular wilt disease on hundreds of plant species, including numerous crops^14,15^. *V. dahliae* survives in the soil through persistent resting structures called microsclerotia that germinate in response to nutrient-rich exudates released by nearby plant roots^16^. Subsequently, emerging hyphae grow through the soil and rhizosphere towards the roots where the fungus penetrates its hosts. Following root penetration, *V. dahliae* invades the xylem where it produces conidiospores that are spread throughout the vasculature by the sap stream. This systemic colonization causes chlorosis and necrosis of plant tissues, which is followed by plant senescence. *V. dahliae* then enters a saprophytic phase, emerges from the vasculature and colonizes the dead plant material where it produces new microsclerotia that are eventually released into the soil upon tissue decomposition.

Using comparative population genomics, we previously identified the *V. dahliae*-secreted small cysteine-rich effector protein Ave1 that is recognized as an avirulence determinant by tomato plants that carry the corresponding Ve1 immune receptor^17^. However, on host plants lacking Ve1, VdAve1 acts as a virulence effector that promotes fungal colonization and disease development^17^. Interestingly, VdAve1 is homologous to plant natriuretic peptides (PNPs) that have been identified in numerous plant species, suggesting that *VdAve1* was acquired from plants through horizontal gene transfer^17^. Whereas several of the plant PNPs were shown to act in plant homeostasis and (a)biotic stress responses^18,19^, the mode of action of VdAve1 to contribute to fungal virulence has remained unknown.

Intriguingly, unlike most pathogen effector genes characterized to date, *VdAve1* is not only highly expressed during host colonization^17,20^, but also during growth *in vitro* and under conditions mimicking soil colonization, suggesting a ubiquitous role throughout the fungal life cycle including life stages outside the host, and thus a role that does not primarily involve targeting host plant physiology (Extended data Fig. 1). Our attempts to purify VdAve1 upon heterologous expression in *Escherichia coli*, to facilitate functional characterization, repeatedly failed due to the formation of inclusion bodies (Extended data Fig. 2a). The inability to obtain soluble protein using heterologous microbial expression systems can be attributed to a multitude of reasons, but is a well-known phenomenon when expressing antimicrobial proteins^21^. Consequently, based on the ubiquitous expression of *VdAve1* by *V. dahliae*, and our inability to purify soluble VdAve1 following expression in *E. coli*, we hypothesized that VdAve1 may possess antimicrobial activity.

To obtain functional VdAve1, inclusion bodies were isolated from *E. coli* cells and denatured using guanidine hydrochloride. Next, VdAve1 was refolded by stepwise dialysis and functionality was confirmed through testing recognition by its immune receptor Ve1 (Extended data Fig. 2b). To assess the potential antimicrobial activity of VdAve1, we developed an *in vitro* system in which we incubated a panel of plant-associated bacteria in tomato xylem fluid, to mimic a natural environment in which VdAve1 is secreted, namely tomato xylem vessels, and monitored their growth in presence and absence of the protein. Interestingly, VdAve1 selectively inhibited the growth of plant-associated bacteria (Fig. 1a). Whereas growth of all Gram positive bacteria tested, namely *Arthrobacter* sp., *Bacillus subtilis, Staphylococcus xylosus* and *Streptomyces* sp., was strongly inhibited, Gram negative bacteria displayed differential sensitivity to the protein. Intriguingly, this differential sensitivity is not immediately explained by phylogenetic relationships of the tested isolates as even within bacterial orders/families differences are observed. For instance, whereas growth of the burkholderiales species *Acidovorax* is inhibited by VdAve1, growth of a Ralstonia isolate, which belongs the same order, is not. Similarly, treatment of two closely related rhizobiales, *Rhizobium* sp. and *Agrobacterium tumefaciens*, revealed differential sensitivity as VdAve1 affected growth of *Rhizobium* sp., but not of *A. tumefaciens*. Finally, growth of Pseudomonas corrugata and *Serratia* sp. was only slightly altered and unaffected, respectively, while growth of both *Sphingobacterium* sp. and *Sphingomonas mali* was affected upon exposure to VdAve1. Interestingly, growth of the endophytic fungus *Fusarium oxysporum* and the fungal mycoparasite *Trichoderma viride* was not inhibited by VdAve1, suggesting that VdAve1 exerts antibacterial, but not antifungal, activity (Extended data Fig. 3a). These initial observations with divergent, randomly chosen, plant-associated bacteria prompted us to further characterize the antimicrobial activity of VdAve1.

**Fig. 1:**
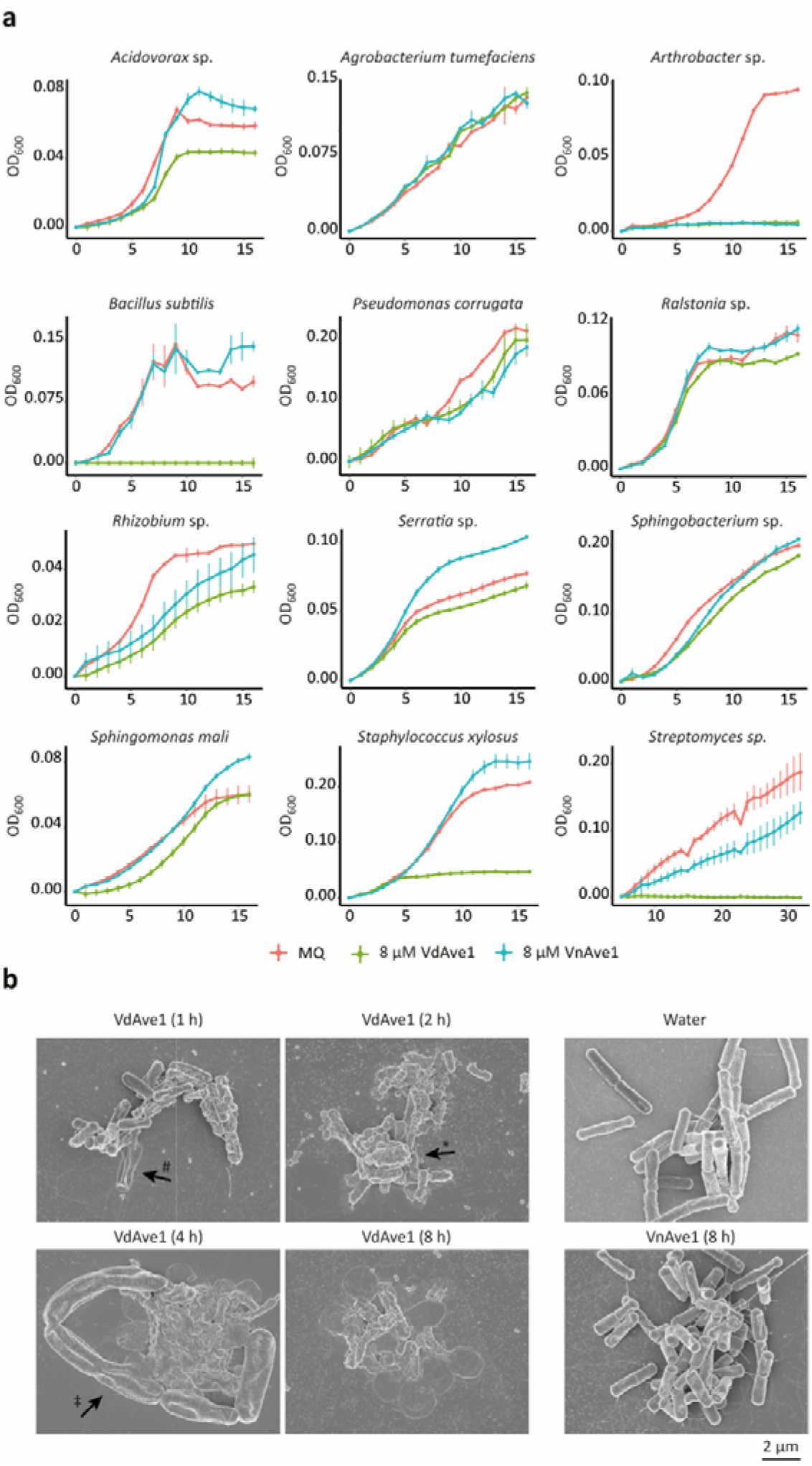
Bactericidal activity of *Verticillium dahliae* effector VdAve1. **a,** VdAve1 selectively inhibits *in vitro* growth of plant-associated bacterial isolates in tomato xylem fluid. The close homolog VnAve1 from *V. nubilum* only inhibits a subset of the bacteria affected by VdAve1 and is generally less effective. **b,** Scanning electron microscopy of *B. subtilis* upon 1, 2, 4 and 8 hours of incubation in tomato xylem fluid showing blebbing (*), swelling (‡) and lysis (#) with 6.5 μM VdAve1 (0.8 × MIC), but not with water or VnAve1.

As a first step in the further characterization of the antimicrobial activity of VdAve1, we aimed to determine whether the effector protein is bacteriostatic or bactericidal by making use of electron microscopy to visualize the effect of protein treatment on bacteria. As a target species the Gram positive *B. subtilis* was chosen, considering its high sensitivity to VdAve1 treatment. By testing a concentration series of the VdAve1 effector protein, the minimum inhibitory concentration (MIC) was determined at 8 μM (Extended data Fig. 3b). However, electron microscopy analysis revealed that sub-MIC concentrations of VdAve1 already induced blebbing and swelling of bacterial cells, followed by lysis and collapse, corresponding with bactericidal activity (Fig. 1b).

To investigate whether the antimicrobial activity that is displayed by VdAve1 is more widely conserved among its homologs, we tested the only homolog that occurs in one of the sister species of the *Verticillium* genus, namely VnAve1, from the non-pathogenic species *V. nubilum* that displays 90% amino acid identity (Extended data Fig. 3c). Interestingly, also this homolog displays antimicrobial activity, albeit that it only inhibits a subset of the bacteria affected by VdAve1, and does not cause *B. subtilis* lysis (Fig. 1). Thus, the 13 amino acid polymorphisms between the two Ave1 homologs are responsible for differences in the activity spectrum. To investigate whether the antimicrobial activity also occurs among plant homologs, or is confined to microbial homologs and involves neofunctionalization after horizontal transfer, the more distant homolog AtPNP-A from *Arabidopsis thaliana* was tested as well. Intriguingly, AtPNP-A completely arrests *B. subtilis* growth (Extended data Fig. 3c,d). Collectively, these findings demonstrate that various Ave1 homologs possess antimicrobial activity, yet with divergent activity spectra, and suggest that the antimicrobial activity of VdAve1 did not result from neofunctionalization following horizontal gene transfer.

Based on the strong but selective bactericidal activity of VdAve1 *in vitro*, we hypothesized that *V. dahliae* exploits its effector protein to affect host microbiome compositions through the suppression of other microbes. Therefore, to determine the biological relevance of the observed bactericidal activity, we performed bacterial community analysis based on 16S ribosomal DNA profiling of tomato and cotton root microbiomes following infection with wild-type *V. dahliae* or a VdAve1 deletion mutant. Importantly, root microbiome compositions were determined during early *V. dahliae* infection stages, namely at ten days post inoculation when the fungus has just entered xylem vessels and initiated systemic spreading, to minimize indirect shifts in microbial compositions that result from severe disease symptomatology, rather than from direct shifts due to the presence of the effector protein. We did not observe major shifts in overall composition of bacterial phyla (Extended data Fig. 4a) or total microbial diversity (α-diversity) (Extended data Fig. 4b) upon *V. dahliae* colonization of tomato and cotton. However, principal coordinate analysis based on Bray-Curtis dissimilarities (β-diversity) revealed a clear separation of root microbiomes (Fig. 2a) (PERMANOVA, p<0.01 for both tomato and cotton). Importantly, the extent of *V. dahliae* colonization does not seem to determine the separation, as clustering of *V. dahliae* genotypes occurs in cotton although *VdAve1* deletion hardly affects fungal virulence on this host plant (Fig. 2a). Thus, as anticipated based on the potent, yet selective, antimicrobial activity, VdAve1 secretion by *V. dahliae* sophistically alters root microbiome compositions. Arguably, based on the sophisticated effects, a full and detailed characterization of microbiome composition changes requires large sample sizes and abundant numbers of repeats. However, strikingly, despite the relatively small sample size of our 16S rDNA profiling, pairwise bacterial order comparisons upon colonization by wild-type *V. dahliae* and the *VdAve1* deletion mutant revealed differential abundances of Sphingomonadales, Bdellovibrionales and Ktedonobacterales for tomato. (Fig. 2b) (Extended data Table 1). The finding that Sphingomonadales are repressed in the presence of VdAve1 suggests that this taxon is the most sensitive to VdAve1 activity. A similar comparison for cotton did not immediately reveal any differentially abundant orders, but agglomeration of amplicon sequence variants (ASVs) based on phylogenetic relatedness (patristic distance<0.1) revealed eight differentially abundant taxa, including a taxon of the Sphingomonadaceae family (Fig. 2b) (Extended data Table 2). Interestingly, although this taxon only represents a small proportion of all Sphingomonadaceae in the cotton root microbiomes, it is exclusively and consistently found in the microbiomes of roots infected by the *VdAve1* deletion mutant, and completely absent upon infection with wild-type *V. dahliae*, again pointing towards the particular sensitivity of this taxon towards VdAve1. Moreover, pairwise comparisons following the combination of tomato and cotton samples based on infection by the different *V. dahliae* genotypes, to identify differentially abundant bacterial orders that potentially remained unnoticed due to the limited sample size, again only revealed differential abundance of Sphingomonadales (p<0.01; Extended data Fig. 4c)(Extended data Table 3).

**Fig. 2:**
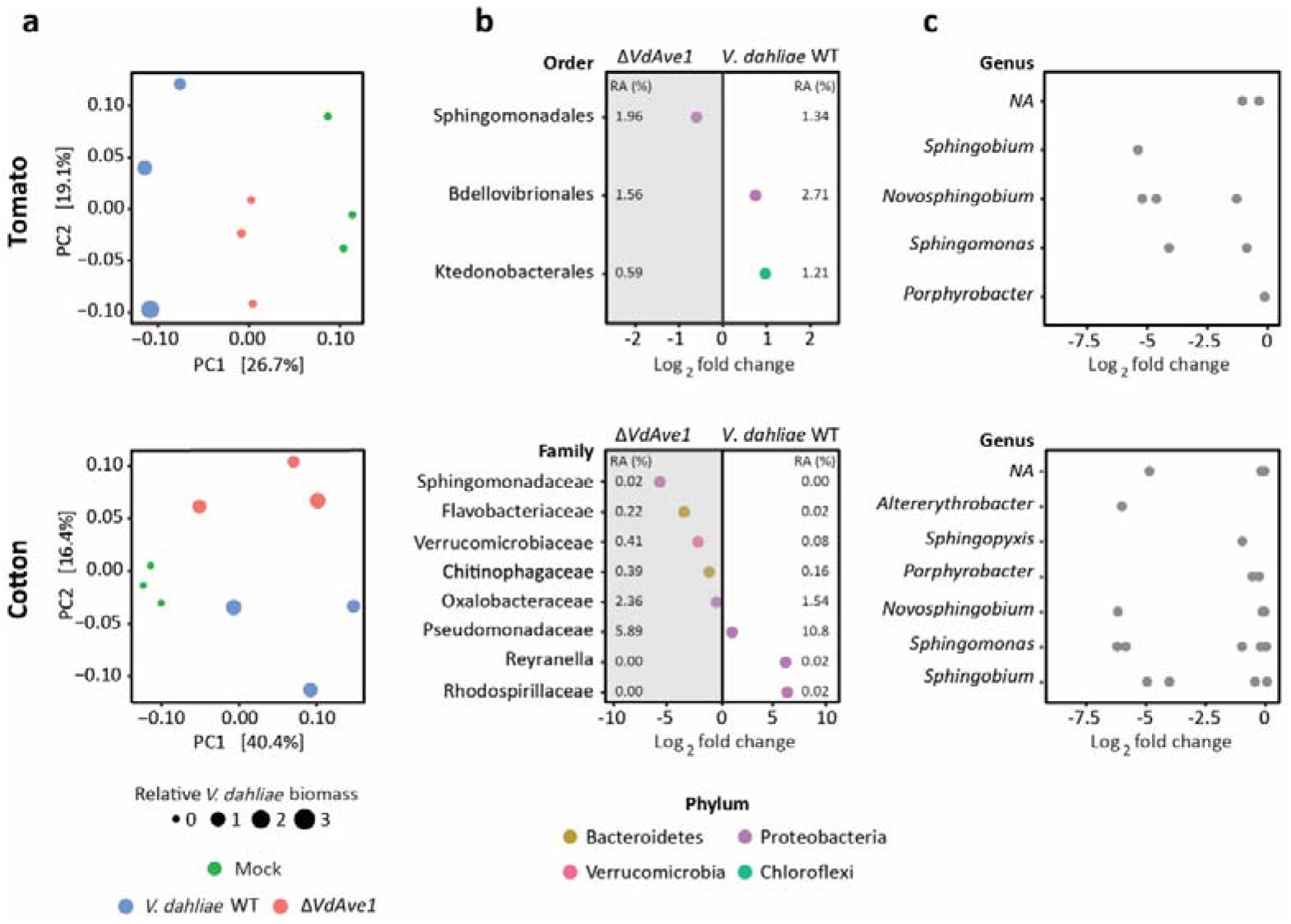
*Verticillium dahliae* VdAve1 impacts root microbiomes. **a,** Principal coordinate analysis based on Bray-Curtis dissimilarities reveals separation of root microbiome compositions ten days after inoculation with wild-type *V. dahliae* and a *VdAve1* deletion mutant (PERMANOVA, p<0.01). **b,** Differential abundance analysis of bacterial orders (tomato) and upon agglomeration of amplicon sequence variants (patristic distance<0.1) (cotton) through pairwise comparison between root microbiomes colonized by wild-type *V. dahliae* and a *VdAve1* deletion mutant (Wald test, p<0.01). The average relative abundance (RA) of the differentially abundant taxa is indicated as a percentage of the total bacterial community in the corresponding root microbiome. **c,** Sphingomonads (*Sphingomonas, Novosphingobium, Sphingopyxis*, and *Sphingobium*) are repressed by *VdAve1*. Dots represent single amplicon sequence variants with increased abundance (average of 3 samples) in root microbiomes upon colonization by the *VdAve1* deletion mutant when compared with wild-type *V. dahliae*.

Given the fact that secretion of VdAve1 by *V. dahliae* during colonization of both tomato and cotton leads to a reduction of Sphingomonadales in the corresponding root microbiomes, we anticipated a broad efficacy of VdAve1 on bacteria within this order. Therefore, to identify Sphingomonadales genera that are most sensitive to VdAve1, we identified ASVs with increased average relative abundance in the microbiomes with the *VdAve1* deletion mutant when compared with wild-type *V. dahliae*, revealing *Sphingomonas, Novosphingobium, Sphingopyxis* and *Sphingobium* that are commonly referred to as Sphingomonads (Fig. 2c)^22,23^. To confirm that the reduced Sphingomonad abundance during *V. dahliae* colonization is a direct consequence of VdAve1 activity, we tested the sensitivity of a panel of plant-associated Sphingomonads to VdAve1 *in vitro*^24,25^. In accordance with the previously observed effect on *S. mali* (Fig. 1a), treatment with VdAve1 was found to also inhibit growth of *Sphingobium, Novosphingobium, Sphingopyxis* and two other *Sphingomonas* species (Fig. 3a), indicating a broad sensitivity among the Sphingomonads. Given the selective efficacy of VdAve1 and the strong effect on Sphingomonads in the tomato and cotton microbiomes, we hypothesized that these bacteria may act as antagonists and negatively affect *V. dahliae* growth in the absence of VdAve1. Indeed, co-cultivation of *V. dahliae* with *Novosphingobium* sp. A and *S. macrogoltabida* resulted in reduced fungal biomass for the VdAve1 deletion mutant, when compared with the *V. dahliae* wild-type that secretes *VdAve1* under these conditions, revealing that Sphingomonads comprise antagonists of *V. dahliae*, and explaining the importance of their inhibition by VdAve1 (Fig. 3b). Accordingly, and in line with previously described observations of plant protective activities of Sphingomonad strains^24^, pre-treatment of surface-sterilized tomato seeds with *S. macrogoltabida* negatively affected *Verticillium* wilt disease development as confirmed through biomass quantification of wild-type *V. dahliae* in the presence and the absence of the bacterium (Fig. 3c,d) (Extended data Fig. 5). Importantly, quantification of *S. macrogoltabida* in the presence of wild-type *V. dahliae* and the *VdAve1* deletion mutant using 16S rDNA profiling and real-time PCR revealed that VdAve1 secretion significantly impacts *S. macrogoltabida* proliferation to counter its protective effect (Fig. 3e-g). Notably, this observation is not an indirect effect of differential host colonization by wild-type *V. dahliae* and the *VdAve1* deletion mutant, as selection of tomato plants with equal levels of *V. dahliae* biomass (Fig. 3d, data points highlighted in red), reveals similarly impaired *S. macrogoltabida* proliferation in the presence of VdAve1 (Fig. 3g). Thus, these data underpin the hypothesis that *V. dahliae* secretes the VdAve1 effector to target antagonistic bacteria, including Sphingomonadales, during host colonization.

**Fig. 3:**
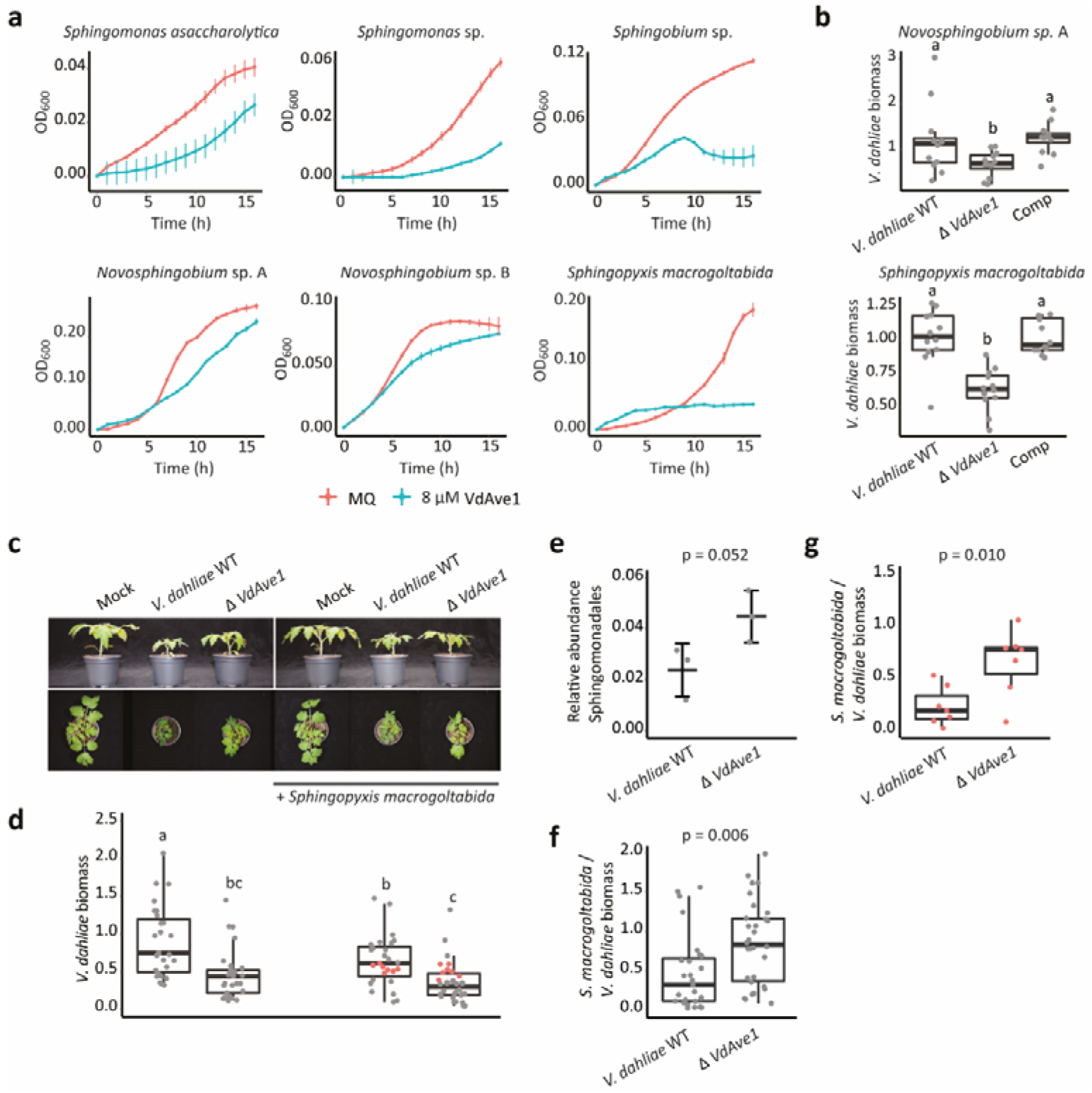
*Verticillium dahliae* VdAve1 affects antagonistic Sphingomonads. **a,** Sphingomonads are inhibited by VdAve1 in tomato xylem fluid. **b,** VdAve1 supports *V. dahliae* growth in the presence of Sphingomonads. Biomass of wild-type *V. dahliae* (WT) and the *VdAve1* deletion (*ΔVdAve1*) and complementation (Comp) mutants was quantified following 48 hours of co-cultivation with Sphingomonads in 0.5x MS medium (N=12). Letters represent significant differences (one-way ANOVA and Tukey’s post-hoc test; p<0.05 for *Novosphingobium* sp.; p<0.0001 for *S. macrogoltabida*). **c,** Tomato seed treatment with *S. macrogoltabida* reduces *Verticillium* wilt symptoms (stunting; 14 days post inoculation). **d,** *V. dahliae* biomass in tomato stems determined with real-time PCR. Letters represent significant biomass differences (one-way ANOVA and Tukey’s post-hoc test; p<0.05; N≥27). Each dot, grey or red, indicates the relative *V. dahliae* biomass in a single tomato plant. **e,** Relative abundance of Sphingomonadales according to 16S ribosomal DNA profiling of tomato plants pre-treated with *S. macrogoltabida* and infected with wild-type *V. dahliae* or the *VdAve1* deletion mutant (unpaired student’s t-test; N=3). **f,** Relative *Sphingopyxis* biomass in all pre-treated tomato plants infected with wild-type *V. dahliae* or the *VdAve1* deletion mutant, indicated by the grey and red dots in Fig. 3d combined, as quantified by real-time PCR (unpaired student’s t-test; N≥27). **g,** Relative *Sphingopyxis* biomass in pre-treated tomato plants colonized by similar amounts of wild-type *V. dahliae* or the *VdAve1* deletion mutant, indicated by the red dots in Fig. 3d, as quantified by real-time PCR (unpaired student’s t-test; N=7).

Our observation that *V. dahliae* secretes VdAve1 to suppress microbial competitors in the microbiomes of its hosts, prompted us to speculate about additional *V. dahliae* effector proteins involved in microbiome manipulation. Based on our findings that canonical effector genes such as *VdAve1* can also be expressed outside the host, we hypothesized that *V. dahliae* also secretes effectors that aid in microbial competition in the soil. Therefore, to query for the occurrence of additional effectors that act in microbiome manipulation, the predicted secretome of *V. dahliae* strain JR2^26^ was probed for structural homologs of known antimicrobial proteins (AMPs), revealing 10 candidates (Extended data Table 4). The majority of the identified effectors share typical characteristics with canonical host-targeting effector proteins, such as being small and rich in cysteines. However, based on previously performed RNA sequencing experiments, no expression of any of these candidates could be monitored during colonization of *Arabidopsis thaliana, Nicotiana benthamiana* or cotton plants (Extended data Fig. 6)^17,20,27,28^. Additionally, *in vitro* cultivation of *V. dahliae* in the presence of *E. coli*, *B. subtilis* or *T. viride*, or of peptidoglycan to mimic bacterial encounter, did not lead to induction of any of the effector candidate genes (Extended data Fig. 6). Consequently, we hypothesized that these genes require other environmental triggers to be induced. Indeed, growth in soil extract consistently induced expression of candidate *VdAMP2* (Fig. 4a) that shares structural homology (confidence >90%) with amphipathic β-hairpins of aerolysin-type β-pore forming toxins (β-PFTs) (Extended data Fig. 7)^29^.

**Fig. 4:**
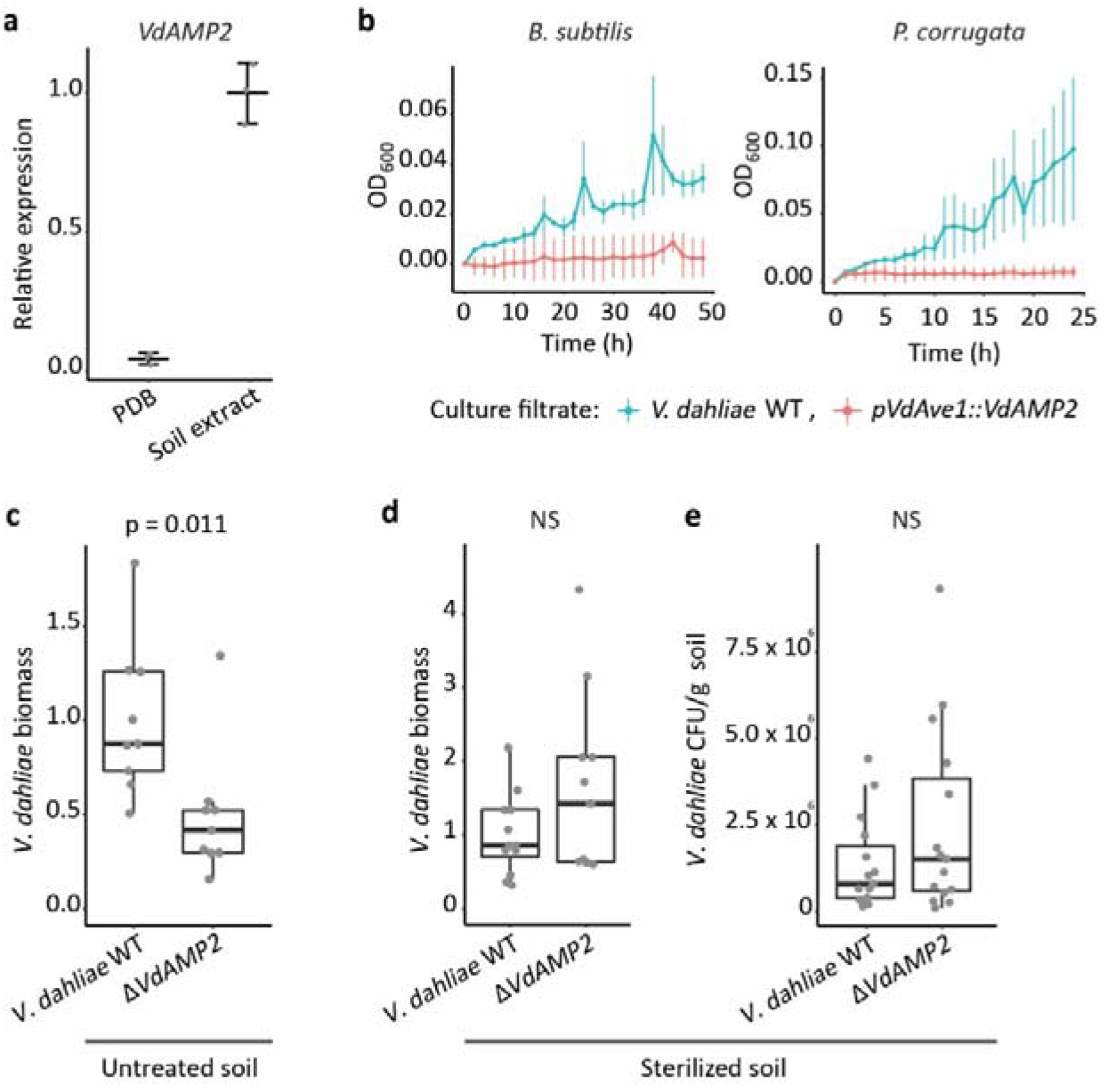
VdAMP2 contributes to *Verticillium dahliae* soil colonization. **a,** *V. dahliae VdAMP2* is induced after five days of cultivation in soil extract but not in potato dextrose broth (PDB). **b,** Growth of *B. subtilis* and *P. corrugata* in filter-sterilized culture filtrates from wild-type *V. dahliae* and the *VdAMP2* expression transformant grown in liquid 0.2× PDB + 0.5× MS medium. **c,** *VdAMP2* contributes to soil colonization. *V. dahliae* biomass in soil samples was determined by real-time PCR seven days after inoculation with wild-type *V. dahliae* (WT) and the *VdAMP2* deletion mutant (Δ*VdAMP2*) (unpaired student’s t-test; N=9). **d,e** VdAMP2 does not contribute to colonization in sterile soil. Experiment as shown in c in sterile soil. *V. dahliae* biomass was quantified with real-time PCR (N=9), **d,** and by colony forming unit counts per gram of soil (N=15), **e.**

To test for potential antimicrobial activity of VdAMP2, we attempted heterologous production of the effector protein. However, since production in *E. coli* and *Pichia pastoris* repeatedly failed, production in *V. dahliae* under control of the *VdAve1* promoter was pursued, resulting in high levels of VdAMP2 expression *in vitro* (Extended data Fig. 8a-c). Interestingly, proliferation of *B. subtilis* and of *P. corrugata* (Fig. 4b), but not of *F. oxysporum* and of *T. viride* (Extended data Fig. 9), was affected by filter-sterilized culture filtrate of the *VdAMP2* expression transformant when compared with that of wild-type *V. dahliae*, suggesting that VdAMP2 exerts only antibacterial activity, like VdAve1 albeit with a different activity spectrum. Soil colonization assays using wild-type *V. dahliae* and a *VdAMP2* deletion mutant (Extended data Fig. 8d-f) demonstrated that VdAMP2 contributes to *V. dahliae* fitness in the soil as measured by biomass accumulation (Fig. 4c). Importantly, since this fitness contribution is not observed in sterilized soil, we conclude that VdAMP2 contributes to *V. dahliae* fitness through its efficacy in microbial competition (Fig. 4d,e). As can be anticipated, the positive effect of VdAMP2 on biomass accumulation in the soil is reflected in disease development when plants are grown on this soil (Extended data Fig. 10), demonstrating that VdAMP2 positively contributes to virulence of *V. dahliae* in an indirect manner.

In conclusion, in this study we have demonstrated that *V. dahliae* employs effector proteins that contribute to niche colonization, during host-associated as well as during soil-dwelling stages, through the selective manipulation of local microbiomes. A wide array of microbially-secreted molecules has previously been described to fulfill crucial functions in intermicrobial competition, including hydrolytic enzymes, secondary metabolites and antimicrobial proteins. Some Gram-negative bacteria even employ a specialized type VI secretion system (T6SS) to translocate antimicrobial proteins into their microbial competitors^30^. In this manner, *Vibrio cholerae*, the causal agent of cholera, employs its T6SS to target members of the host commensal microbiota and hereby promotes colonization of the gut^31^. Similarly, the T6SS effector Hyde1 of the phytopathogen *Acidovorax citrulli* targets plant-associated bacteria *in vitro* and was speculated to play a role in microbial competition *in planta*^7^. This T6SS is analogous to the type III secretion system (T3SS) of Gram negative bacteria that acts as a needle-like structure to directly inject effector proteins into host cells to promote disease^32^. Similar secretion machinery intended for host-microbe or microbe-microbe interactions has not been described for fungi and other filamentous microbes, which instead secrete their effector proteins by extracellular deposition. Consequently, effector molecules targeted towards host cells or towards microbial competitors cannot be discriminated based on differential secretion motifs, such as those that determine type III versus type VI secretion in Gram negatives. Here, we have shown that the pool of effectors secreted by a fungal plant pathogen represents a diverse cocktail comprising proteins involved in the manipulation of the host as well as its microbiome. Consequently, the effectors reported here likely only represent a small proportion of a larger subset of the *V. dahliae* effector repertoire that is intended for microbiome manipulation. For instance, similar effectors might be crucial during advanced infection stages to prevent secondary infections by opportunistic microbes when host defenses are impaired. Additionally, effector proteins can be anticipated to facilitate the survival of the *V. dahliae* resting structures that persist in the microbe-rich soil for years^33^. After all, possibly, fungal effectors with host microbiome-manipulating capacity initially evolved to limit bacterial growth in soil, as the advent of fungi on earth preceded land plant evolution and fungi initially likely co-evolved with bacteria in soil to compete for organic carbon. The discovery of further molecules for microbiome manipulation secreted by *V. dahliae* and other microbes, and unravelling of underlying modes of action, may ultimately lead to the development of novel antibiotics.

## Materials and methods

All experiments have been repeated at least three times.

### Xylem fluid isolation

Tomato plants (*Solanum lycopersicum* cv. Moneymaker) were grown under controlled greenhouse conditions as described previously^34^. The stems of six-week-old plants were cut to allow oozing of the xylem fluid, which was collected on ice with a vacuum pump. The collected xylem fluid was centrifuged for 10 minutes at 20000 × *g* and filter-sterilized using a 0.2 μm filter (Sarstedt, Nümbrecht, Germany). The sterilized xylem fluid was stored at −20°C until use.

### Soil extract preparation

To prepare soil extract, 100 grams of dry potting soil (Lentse potgrond, substraat arabidopsis, Lentse Potgrond BV, Katwijk, the Netherlands) was mixed with 500 mL of demineralized water and autoclaved for 15 minutes at 121°C. Soil particles were pelleted through centrifugation and the supernatant was collected and stored at −20°C until use.

### Gene expression analysis

Total RNA of *V. dahliae* strain JR2 was isolated from tomato roots seven days after root dip inoculation and following five days of *in vitro* growth in soil extract and potato dextrose broth (PDB) using the Maxwell^®^ 16 LEV Plant RNA Kit (Promega, Madison, USA). Real-time PCR was performed as described previously^17^ to determine the expression of effector genes relative to *VdGAPDH* with primer pairs as shown in Extended data Table 6.

### Production and purification of recombinant effector proteins

The sequences encoding mature VdAve1 and VnAve1 were cloned into pET-15b with an N-terminal His_6_ tag sequence (Novagen, Madison, WI, USA) (primer sequences, see Extended data Table 6). The resulting expression vectors were confirmed by sequencing and used to transform *E. coli* strain BL21. For heterologous protein production, BL21 cells were grown in 1 × YT liquid medium at 37°C with constant shaking at 200 rpm. Protein production was induced with 1 mM IPTG final concentration when cultures reached an OD_600_=2 to ensure maximum yields. Following 2 hours of protein production, the bacterial cells were pelleted and snap-frozen in liquid nitrogen and then washed with 100 mM NaCl, 1 mM EDTA, and 10 mM Tris at pH 8.5. Cells were disrupted by stirring for 1 hour in lysis buffer (100 mM Tris, 150 mM NaCl, 10% glycerol, 6 mg/mL lysozyme (Sigma, St. Louis, MO, USA), 2 mg/mL deoxycholic acid, 0.06 mg/mL DNaseI, protease inhibitor cocktail (Roche, Mannheim, Germany)) at 4°C. Soluble and insoluble fractions were separated by centrifuging at 20,000 × *g* for 10 min. The insoluble protein pellets were washed with 10 mL 1 M guanidine hydrochloride (GnHCl), 10 mM Tris at pH 8.0 and then denatured in 10 mL 6 M GnHCl, 10 mM β-mercaptoethanol, 10 mM Tris at pH 8.0. Samples were incubated for 1 hour at room temperature. Non-denatured debris was pelleted by centrifuging at 20,000 × *g* for 10 min and discarded. Denaturation was allowed to continue for additional 3-4 hours. Proteins were purified under denaturing conditions by metal affinity chromatography using a column packed with 50% His60 Ni^2+^ Superflow Resin (Clontech, Mountain View, CA, USA). The purified effector proteins were dialysed (Spectra/Por^®^3 Dialysis Membrane, MWCO= 3.5 kDa) step-wise against 20 volumes of 0.25 M ammonium sulfate, 0.1 M BisTris, 10 mM reduced glutathione, 2 mM oxidized glutathione, pH 5.5 with decreasing GnHCl concentrations for refolding. Each dialysis step was allowed to proceed for at least 24 hours. Finally, proteins were dialysed against demineralized water. Final concentrations were determined using the BioRad Protein Assay (BioRad, Veenendaal, The Netherlands).

Functionality of refolded VdAve1 was confirmed through recognition by the corresponding tomato immune receptor Ve1. To this end, an overnight culture of *A. tumefaciens* strain GV3101 carrying the pSOL2092:Ve1 construct^35^ was harvested by centrifugation and re-suspended to OD_600_ =2 in MMA (2% sucrose, 0.5% Murashige & Skoog salts (Duchefa Biochemie, Haarlem, The Netherlands), 10 mM MES, 200 μM acetosyringone, pH 5.6) and infiltrated in the leaves of 5-week-old *N. tabacum* (cv. Petite Havana SR1) plants. After 24 hours, 10 μM of purified and refolded 6xHis-VdAve1 was infiltrated in leaf areas expressing Ve1. Photos were taken three days post infiltration of the effector protein.

### Generation of *V. dahliae* mutants

To generate the *VdAMP2* effector deletion construct, *VdAMP2* flanking sequences were amplified using the primers listed in Extended data Table 6 and cloned into pRF-HU2^36^. To allow expression of *VdAMP2* under control of the VdAve1 promoter, the coding sequence of *VdAMP2* was amplified and cloned into pFBT005. All constructs were transformed into *A. tumefaciens* strain AGL1 for *V. dahliae* transformation as described previously^37^.

### *V. dahliae* culture filtrates

Conidiospores of *V. dahliae* strain JR2 and the *VdAMP2* expression transformant were harvested from potato dextrose agar (PDA) and diluted to a final concentration of 10^4^ conidiospores/mL in 20 mL of 0.2× PDB supplemented + 0.5× Murashige & Skoog medium (Duchefa, Haarlem, The Netherlands). Following four days of incubation at 22°C and 120 rpm, the fungal biomass was pelleted and the remaining supernatants were filter sterilized and stored at −20°C until use.

### Bacterial isolates

Bacterial strains *Bacillus subtilis* AC95, *Staphylococcus xylosus* M3, *Pseudomonas corrugata* C26, *Streptomyces* sp. NE-P-8 and *Ralstonia* sp. M21 were obtained from our in house endophyte culture collection. Strains used in this study were all isolated from the xylem vessels of tomato cultivars from commercial greenhouses, both from stem and leaf sections. All strains were identified based on their 16S rRNA gene sequence using the primers 27F and 1492R (Extended data Table 6). 16S amplicons were sequenced by Sanger sequencing at Eurofins (Mix2Seq). The partial 16S rRNA gene sequences obtained were evaluated against the 16S ribosomal DNA sequence (Bacteria and Archaea) database from NCBI. Bacterial strains *Acidovorax* sp. (Leaf 73), *Arthrobacter* sp. (Leaf 69), *Rhizobium* sp. (Leaf 167), *Serratia* sp. (Leaf 50), *Sphingomonas* sp. (Leaf 198), *Sphingobium* sp. (Leaf 26) and *Novosphingobium* sp. B (Leaf 2) were obtained from the At-SPHERE collection^25^. Bacterial strains *S. mali* (DSM 10565) and *S. asaccharolytica* (DSM 10564) were obtained from the DSMZ culture collection (Braunschweig, Germany). Bacterial strains *Novosphingobium* sp. A (NCCB 100261), *S. macrogoltabida* (NCCB 95163), and *Sphingobacterium* sp. (NCCB 100093) were obtained from the Westerdijk Fungal Biodiversity Institute (Utrecht, The Netherlands).

### *In vitro* microbial growth assays

Bacterial isolates were grown on lysogeny broth agar (LBA) or tryptone soya agar (TSA) at 28°C. Single colonies were selected and grown overnight at 28°C while shaking at 200 rpm. Overnight cultures were resuspended to OD_600_=0.05 in xylem fluid supplemented with purified effector proteins or diluted using culture filtrates to OD_600_=0.1. Additionally, *F. oxysporum* and *T. viride* spores were harvested from a PDA plate and suspended in xylem fluid supplemented with purified effector proteins or the *V. dahliae* culture filtrates to a final concentration of 10^4^ spores/mL. 200 μL of the microbial suspensions was aliquoted in clear 96 well flat bottom polystyrene tissue culture plates. Plates were incubated in a CLARIOstar^®^ plate reader (BMG LABTECH, Ortenberg, Germany) at 22°C with double orbital shaking every 15 minutes (10 seconds at 300 rpm). The optical density was measured every 15 minutes at 600 nm.

### Scanning electron microscopy

Samples for scanning electron microscopy were prepared as described previously with slight modifications^38^. In short, *B. subtilis* strain AC95 was grown overnight in LB and resuspended in xylem fluid to an OD_600_=0.05. Purified effector proteins were added to a final concentration of 6.5 μM (= 0.8 × MIC, VdAve1) and bacterial suspensions were incubated for 0, 1, 3 and 7 hours. Next, 20 μL of the bacterial suspensions was transferred to poly-L-lysine coated glass slides (Corning, New York, USA) and incubated for another hour to allow binding of the bacteria. Glass slides were washed using sterile MQ and samples were fixed using 2.5% glutaraldehyde followed by postfixation in 1% osmium tetroxide. Samples were dehydrated using an ethanol dehydration series and subjected to critical point drying using a Leica CPD300 (Leica Mikrosysteme GmbH, Vienna, Austria). Finally, the samples were mounted on stubs, coated with 12nm of tungsten and visualized in a field emission scanning electron microscope (Magellan 400, FEI, Eindhoven, the Netherlands).

### Root microbiome analysis

Tomato and cotton inoculations were performed as described previously^34^. After ten days, plants were carefully uprooted and gently shaken to remove loosely adhering soil from the roots. Next, roots with rhizosphere soil from three tomato or two cotton plants were pooled to form a single biological replicate. Samples were flash-frozen in liquid nitrogen and ground using mortar and pestle. Genomic DNA isolation was performed using the DNeasy PowerSoil Kit (Qiagen, Venlo, The Netherlands). Quality of the DNA samples was checked on a 1.0% agarose gel. Sequence libraries were prepared following amplification of the V4 region of the bacterial 16S rDNA (515F and 806R), and paired ends (250 bp) were sequenced using the HiSeq2500 sequencing platform (Illumina, San Diego, USA) at the Beijing Genome Institute (BGI, Hong Kong, China).

Sequencing data was processed using R version 3.3.2. as described previously^39^. Briefly, amplicon sequence variants (ASVs) were inferred from quality filtered reads (Phred score >30) using the DADA2 method^40^. Taxonomy was assigned using the Ribosomal Database Project training set (RDP, version 16) and mitochondria- and chloroplast-assigned ASVs were removed. Next, ASV frequencies were transformed according to library size to determine relative abundances. The phyloseq package (version 1.22.3) was used to determine α-diversity (Shannon index) and β-diversity (Bray-Curtis dissimilarity) as described previously^33,41^. Differential abundance analysis was performed using the DESeq2 extension within phyloseq^42^. To this end, a parametric model was applied to the data and a negative binomial Wald test was used to test for differential abundance of bacterial taxa with p<0.01 as significance threshold.

### *In vitro* competition assay

Conidiospores of *V. dahliae* strain JR2 and the *VdAve1* deletion and complementation mutants were harvested from a PDA plate using sterile water and diluted to a final concentration of 10^6^ conidiospores/mL in liquid 0.5× MS (Murashige and Skoog) medium (Duchefa, Haarlem, The Netherlands). Next, overnight cultures of *Novosphingobium sp.* and *S. macrogoltabida* were added to the conidiospores to OD_600_=0.05 and 500 μL of the microbial suspensions was aliquoted in clear 12-well flat-bottom polystyrene tissue culture plates. Following 48 hours of incubation at room temperature, the microbial cultures were recovered and genomic DNA was isolated using the SmartExtract - DNA Extraction Kit (Eurogentec, Maastricht, The Netherlands). *V. dahliae* biomass was quantified through real-time PCR using *V. dahliae* specific primers targeting the internal transcribed spacer (ITS) region of the ribosomal DNA (Extended data Table 6).

### *In planta* competition assay

To allow *S. macrogoltabida* colonization of in the absence of other microbes, tomato seeds were incubated for five minutes in 2% sodium hypochlorite to ensure surface sterilization. Next, surface sterilized tomato seeds were washed three times using sterile water and transferred to a sterile Petri dish containing a filter paper pre-moistened with a *S. macrogoltabida* suspension in water (OD_600_=0.05). The tomato seeds were allowed to germinate *in vitro* and eventually transferred to regular potting soil, ten-day-old seedlings were inoculated as described previously ^34^. Tomato stems were collected at 14 days post inoculation (dpi) and lyophilized prior to genomic DNA isolation with a CTAB-based extraction buffer (100 mM Tris-HCl pH 8.0, 20 mM EDTA, 2 M NaCl, 3% CTAB). *V. dahliae* biomass was quantified with real-time PCR on the genomic DNA by targeting the internal transcribed spacer (ITS) region of the ribosomal DNA. The tomato *rubisco* gene was used for sample calibration. *S. macrogoltabida* biomass was quantified using Sphingopyxis specific primers (Extended data Table 6) and normalized using the *V. dahliae* ITS. Additionally, the relative abundance of the Sphingomonadales in three representative samples was determined by 16S ribosomal DNA profiling as described previously.

### Soil colonization assays

Conidiospores of the *V. dahliae* strain JR2 and the mutants were harvested from PDA plate and a total of 10^6^ conidiospores were added to 1 gram of potting soil. Samples were incubated at room temperature in the dark. After one week, DNA was extracted from the soil samples using the DNeasy PowerSoil Kit (QIAGEN, Venlo, The Netherlands). *V. dahliae* biomass was quantified through real-time PCR using *V. dahliae* specific primers targeting the internal transcribed spacer (ITS) region of the ribosomal DNA (Extended data Table 6). Primers targeting a conserved region of the bacterial 16S rRNA gene were used for sample equilibration.

To allow sample calibration when using sterilized potting soil (15 minutes at 121°C), the samples were first mixed in a 1:1 ratio with fresh potting soil prior to DNA extraction. Additionally, after one week of incubation of *V. dahliae* in the sterilized soil, serial dilutions were made and plated onto PDA to quantify colony forming units.

### Disease assays using *V. dahliae* microsclerotia

*V. dahliae* microsclerotia were produced in a sterile moist medium of vermiculite and maize meal as described previously^43^. After four weeks of incubation, the vermiculite/microsclerotia mixture was dried at room temperature. Next, 150 mL of the dried mixture was mixed with 1 L of potting soil (Lentse potgrond, substraat arabidopsis, Lentse Potgrond BV, Katwijk The Netherlands) and Arabidopsis seeds of the Col-0 ecotype were sown at equal distances on top of the mixture. The above-ground parts of the plants were collected at 27 dpi and *V. dahliae* biomass was quantified through real-time PCR using *V. dahliae* specific primers targeting the internal transcribed spacer (ITS) region of the ribosomal DNA. The Arabidopsis rubisco gene was used for sample calibration (Extended data Table 6).

## Acknowledgments

The authors thank M. Giesbers from the Wageningen Electron Microscopy Centre for technical assistance. Work in the laboratory of B.P.H.J.T is supported by the Research Council Earth and Life Sciences (ALW) of the Netherlands Organization of Scientific Research (NWO).

## Author contributions

N.C.S., H.R. and B.P.H.J.T. conceived the project. N.C.S., H.R., G.C.P., J.R.M. and B.P.H.J.T. designed the experiments. N.C.S., H.R., G.C.P., M.R.F., M.F.S. and R.N. carried out the experiments, N.C.S., H.R., G.C.P., M.R.F., M.F.S., J.A.V., R.N., J.R.M. and B.P.H.J.T. analysed the data. N.C.S. and B.P.H.J.T. wrote the manuscript. All authors read and approved the final manuscript.

## Competing interests

Authors declare no competing interests.

## Data and materials availability

The metagenomics data have been deposited in the European Nucleotide Archive (ENA) under accession number PRJEB34281.

