## Supplemental Data for "A plant pathogen utilizes effector proteins for microbiome manipulation"

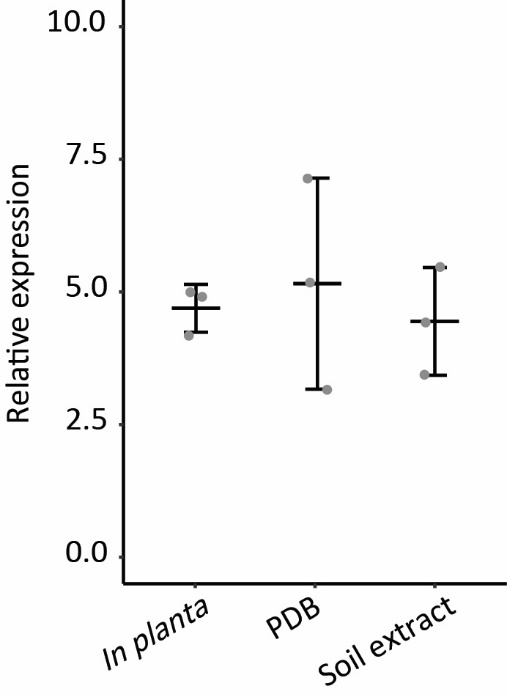

**Extended data Fig. 1: The *VdAve1* effector gene is ubiquitously expressed by *Verticillium dahliae*.** *VdAve1* is among the most highly expressed effector genes *in planta*^17,20,44^. The graph displays expression of *VdAve1* relative to *VdGAPDH* during colonization of tomato roots at 7 days post inoculation, growth in potato dextrose broth (PDB) at 5 days of cultivation, or growth in soil extract at 5 days of cultivation. Error bars show the standard deviation of three biological replicates.

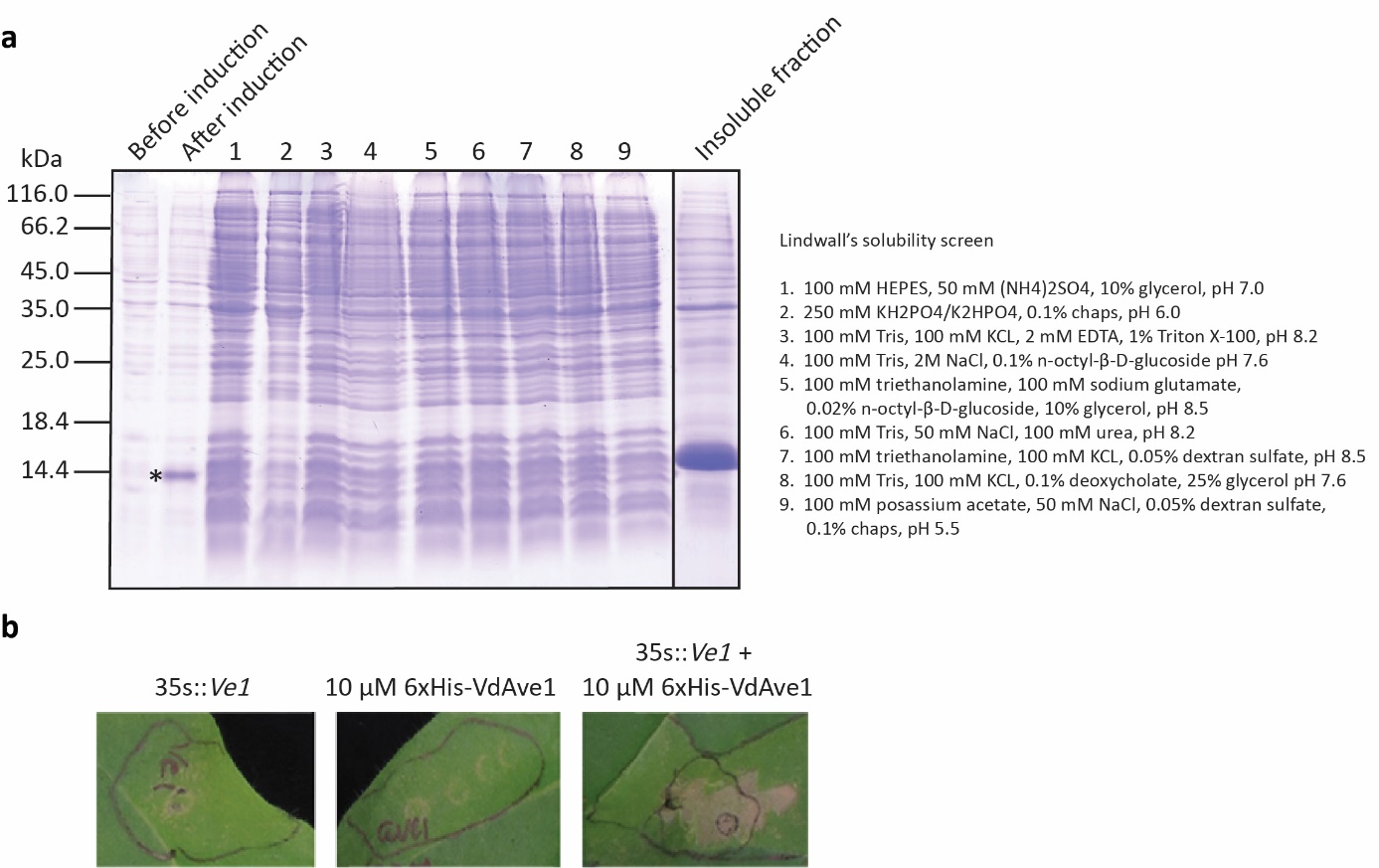

**Extended data Fig. 2: Heterologously produced VdAve1 can be isolated from inclusion bodies. a,** *E. coli* BL21 cells were grown in liquid YT medium and *VdAve1* expression was induced using 1 mM IPTG. Following four hours of protein production at 28^o^C, the presence of VdAve1 was confirmed by boiling the cells in 1% SDS, 2M urea, 1.25% β-mercaptoethanol, 2.5% glycerol, 15 mM Tris, pH 6.8. The band representing VdAve1 is indicated with an asterisk; limited solubility of the protein was detected upon sonication of the cells in the corresponding buffers (lanes 1-9). The presence of VdAve1 in the insoluble protein fractions was confirmed following denaturation of the insoluble proteins, indicating the formation of inclusion bodies. **b,** VdAve1 purified from the insoluble protein fraction under denaturing conditions was refolded by step-wise dialysis. Functionality of the protein was confirmed by infiltration into *N. tabacum* leaf sections overexpressing the corresponding tomato immune receptor Ve1, resulting in a hypersensitive response at three days post infiltration.

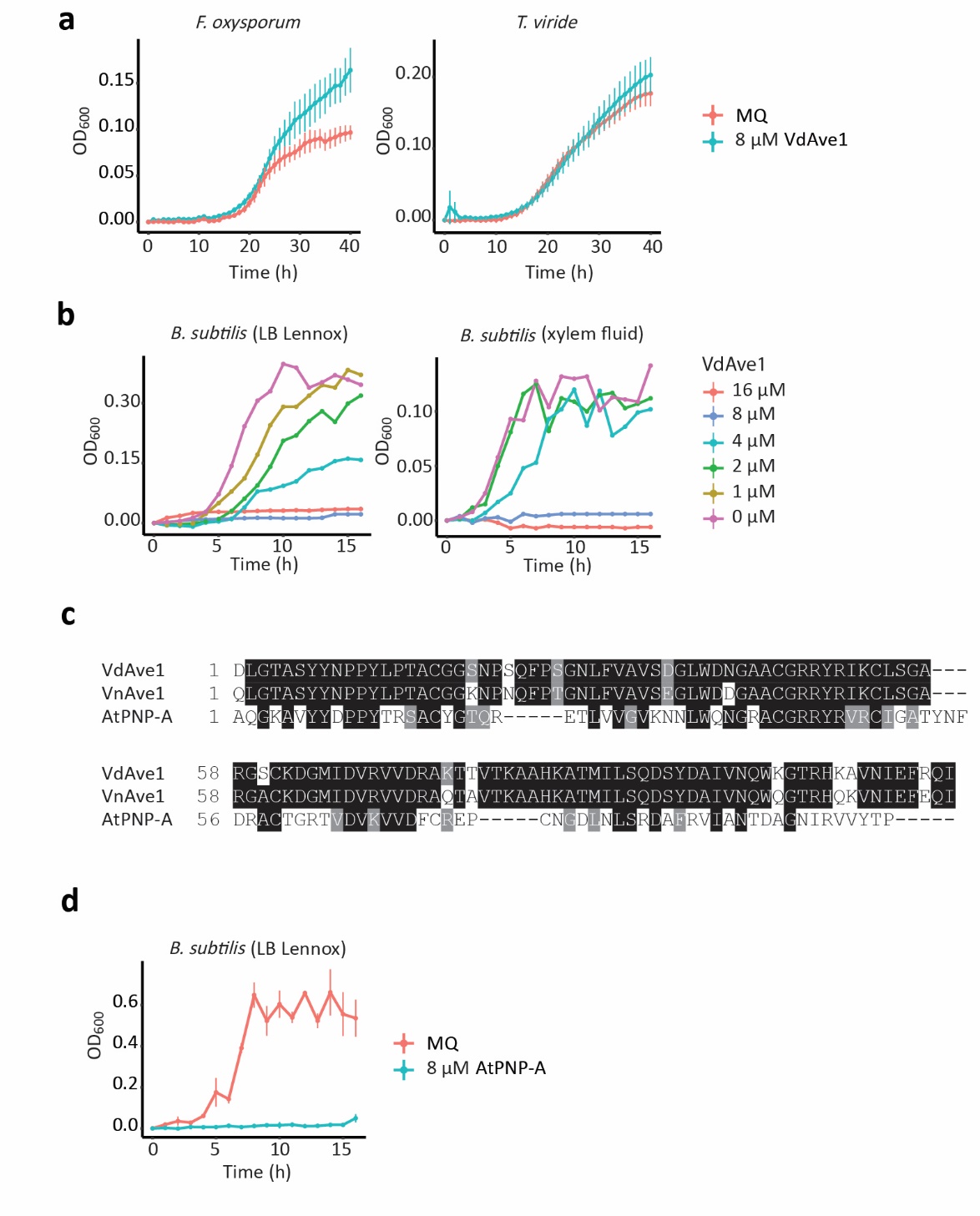

**Extended data Fig. 3: VdAve1 displays antibacterial, but not antifungal, activity. a,** *In vitro* growth of *F. oxysporum* and *T. viride* in tomato xylem fluid is not inhibited by 8 µM VdAve1. **b,** Determination of the minimum inhibitory concentration of VdAve1 on *B. subtilis* upon overnight incubation in LB Lennox and tomato xylem fluid. No growth was detected upon incubation with 8 µM VdAve1 or more. **c,** Multiple sequence alignment of mature VdAve1 with the homologs from *V. nubilum* and *A. thaliana*. **d,** *In vitro* growth of *B. subtilis* is inhibited by 8 µM AtPNP-A.

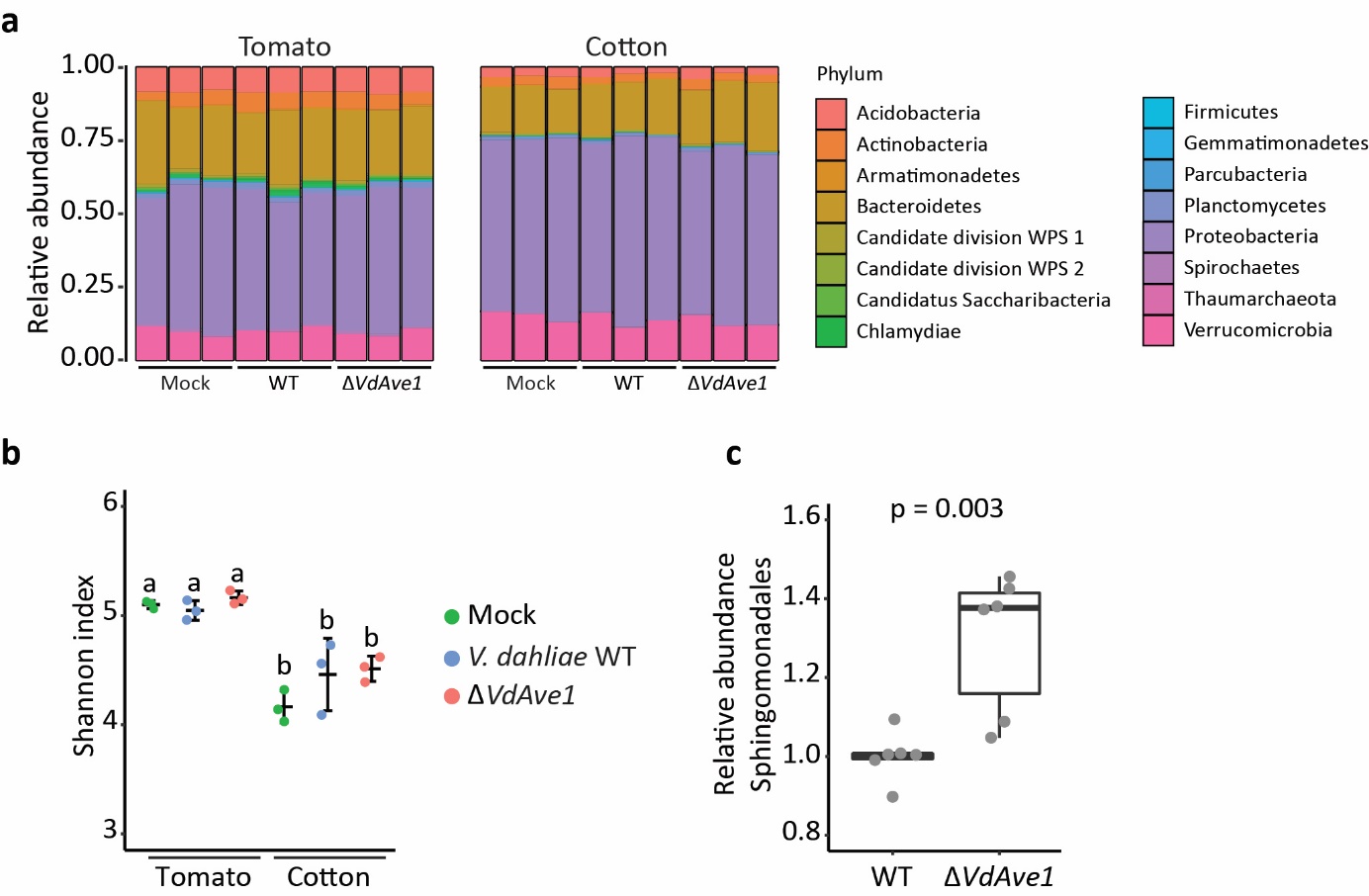

**Extended data Fig. 4: Metagenomic characterization of tomato and cotton root microbiomes upon *Verticillium dahliae* infection. a,** Relative abundance of bacterial phyla in the root microbiomes of tomato and cotton plants ten days after inoculation with wild-type *V. dahliae* (WT) and a *VdAve1* deletion mutant as determined by 16S ribosomal DNA profiling. **b,** *V. dahliae* inoculation does not change α-diversity of host root microbiomes. **c,** Sphingomonadales are significantly enriched in the microbiomes of roots that are colonized by the *VdAve1* deletion mutant. Differential abundance analysis of bacterial orders following combination of tomato and cotton samples based on infection by the different *V. dahliae* genotypes, only revealed the Sphingomonadales as differentially abundant (unpaired student’s t-test, p<0.01; N=6). Relative abundances were normalized against the average relative abundance upon infection of the corresponding host by wild-type *V. dahliae* to correct for host-dependent differences.

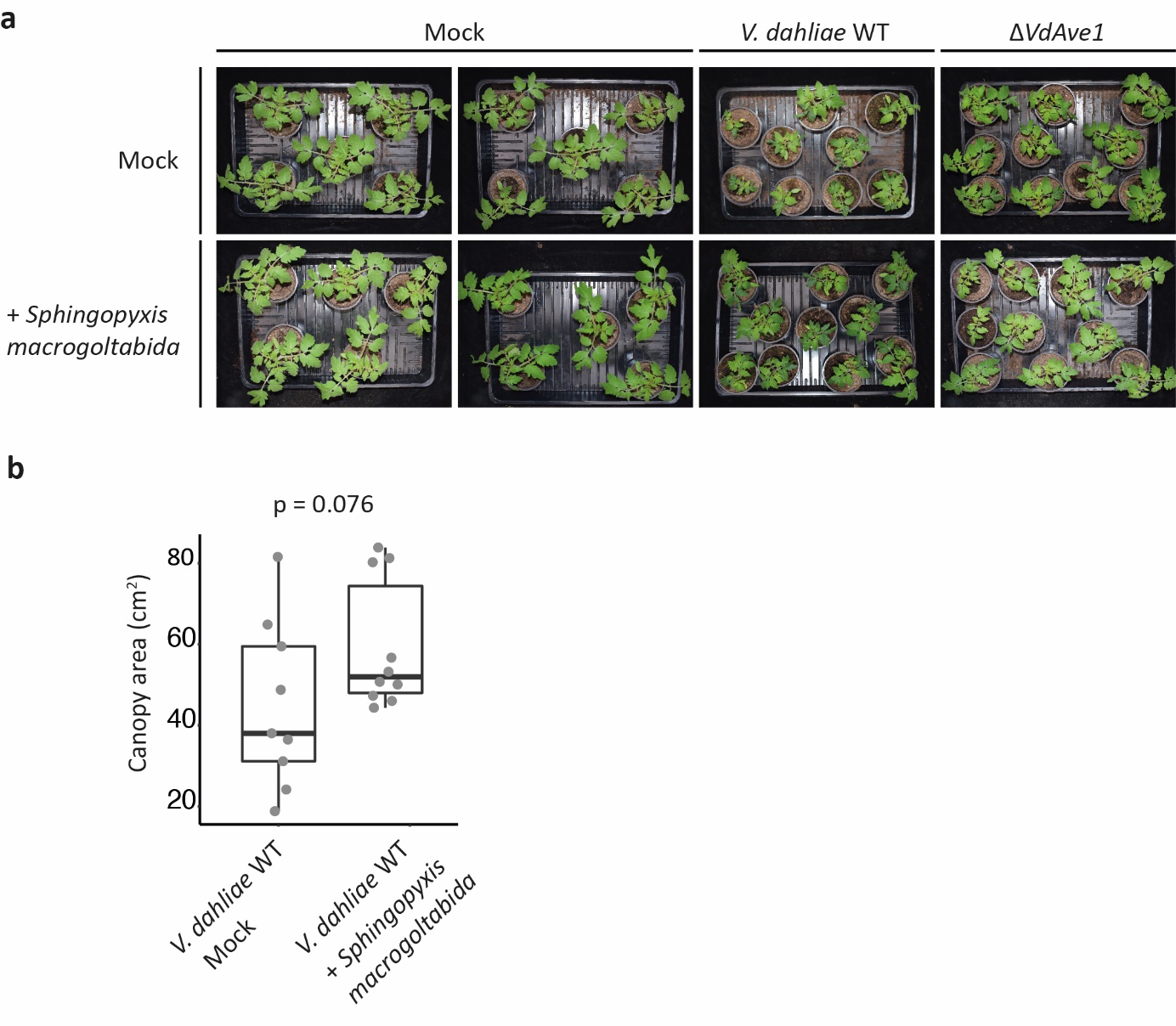

**Extended data Fig. 5: Tomato seed treatment with *S. macrogoltabida* reduces Verticillium wilt symptoms. a,** Phenotypes of tomato plants 14 days post inoculation with wild-type *V. dahliae* or the *VdAve1* deletion mutant. Tomato seeds were surface-sterilized and allowed to germinate *in vitro* in the presence or the absence of *S. macrogoltabida* prior to infection. **b,** Canopy area of mock-treated and *S. macrogoltabida*-treated tomato plants infected by wild-type *V. dahliae* (unpaired student’s t-test; N≥9).

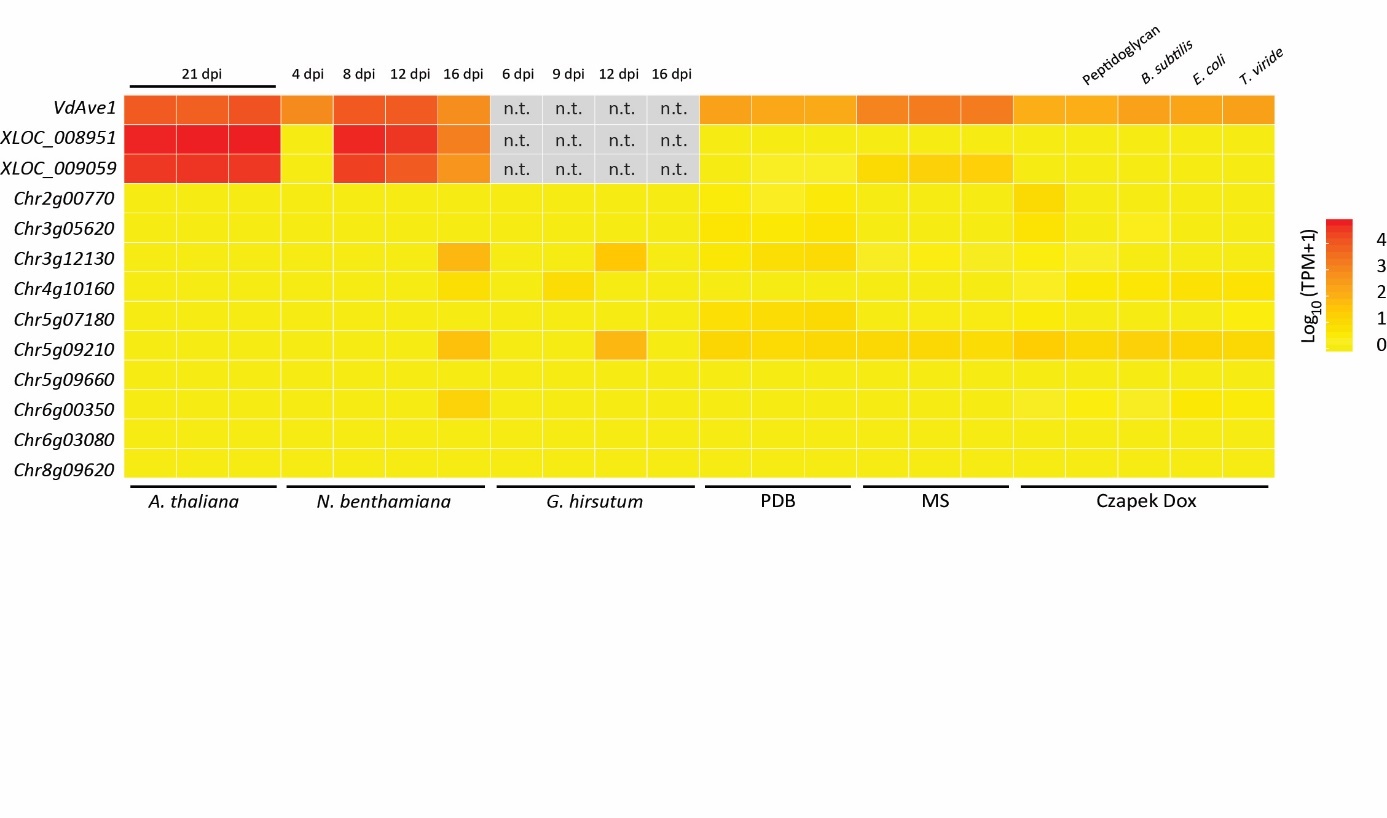

**Extended data Fig. 6: Expression analysis of putative antimicrobial effector gene candidates.** Expression profiles of ten putative antimicrobial effector gene candidates with three previously characterized lineage-specific *in planta*-induced effectors (VdAve1, XLOC_008951, XLOC_009059) during host colonization in *A. thaliana*, *N. benthamiana*, *G. hirsutum* (cotton) and *in vitro* growth in potato dextrose broth (PDB), 0.5x MS (Murashige and Skoog) medium and Czapek Dox in the presence and absence of peptidoglycan, *B. subtilis*, *E. coli* and *T. viride (15, 16, 20, 21)*. The log-scaled color gradient indicates gene expression in Transcripts Per Million (TPM), gray boxes indicate absence of the lineage specific effector genes in *V. dahliae* strain CQ2 that was used for cotton infection. All the other expression profiles were generated using *V. dahliae* strain JR2.

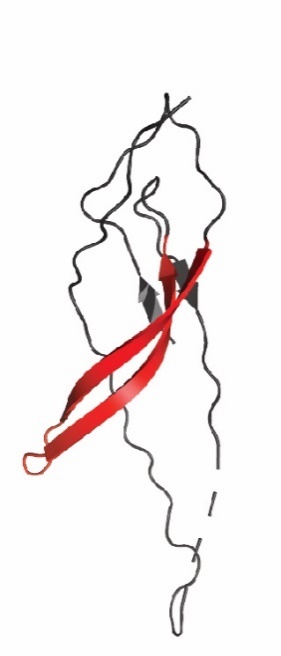

**Extended data Fig. 7: VdAMP2 shares structural homology with the amphipathic β-hairpins of aerolysin-type β-pore forming toxins.** Predicted protein structure of part of VdAMP2 (aa 225-312) with the amphipathic β-hairpin highlighted in red. Protein structure was predicted using Phyre2^45^.

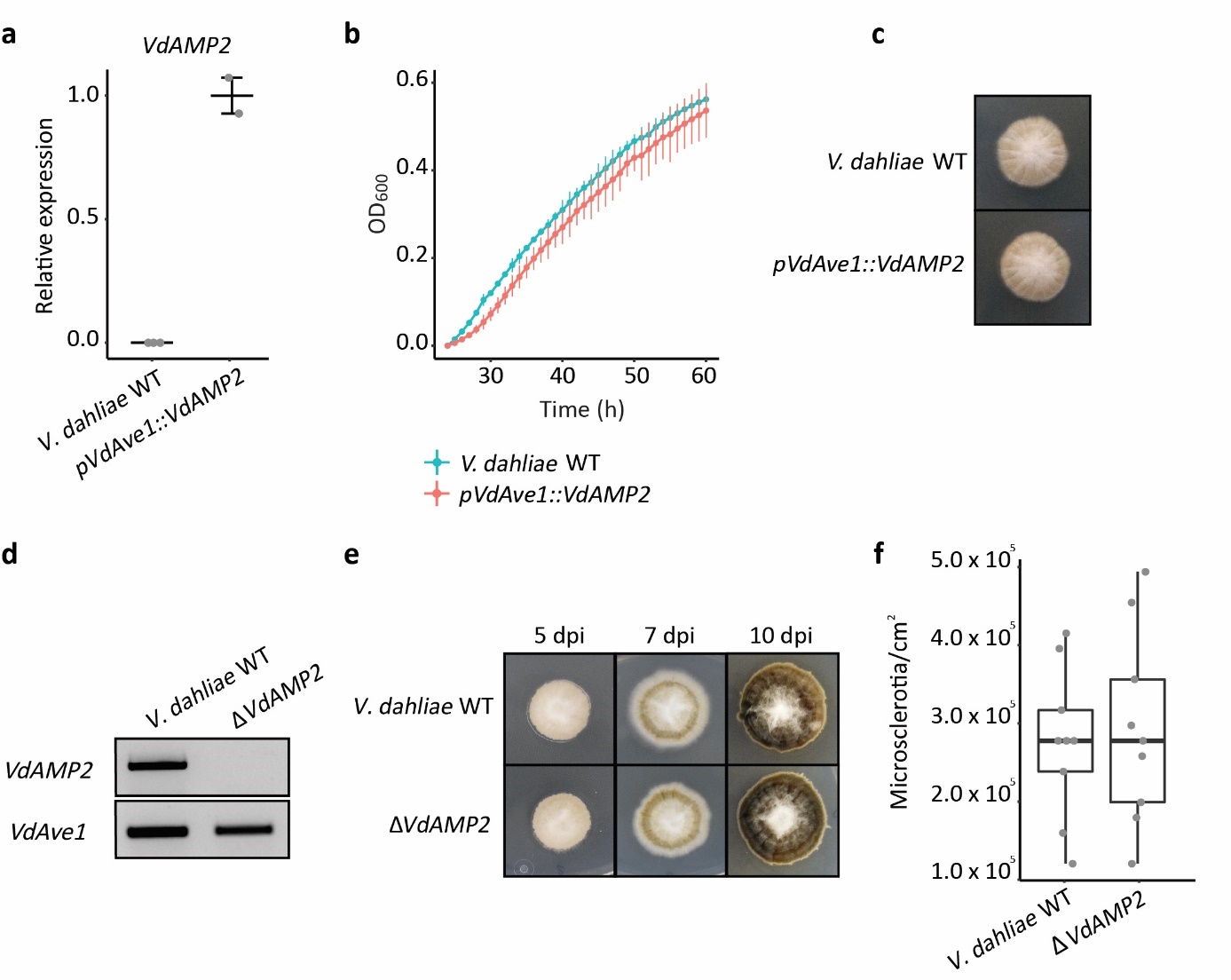

**Extended data Fig. 8: *Verticillium dahliae VdAMP2* mutants. a,** *In vitro* expression of *VdAMP2* in wild-type *V. dahliae* and the *pVdAve1::VdAMP2* mutant after five days of cultivation in liquid 0.5x Murashige & Skoog (MS) medium. **b-c,** *In vitro* expression of *VdAMP2* does not affect *V. dahliae* growth. **b,** Growth of wild-type *V. dahliae* and the *pVdAve1::VdAMP2* mutant in liquid 0.2x potato dextrose broth (PDB) + 0.5x MS. **c,** Morphology of wild-type *V. dahliae* and the *pVdAve1::VdAMP2* mutant after five days of cultivation on potato dextrose agar (PDA). **d,** Deletion of *VdAMP2* was confirmed using PCR on genomic DNA of wild-type *V. dahliae* and the *VdAMP2* deletion mutant (Δ*VdAMP2*), *VdAve1* was used as genomic DNA control. **e,** Morphology of wild-type *V. dahliae* and the *VdAMP2* deletion mutant at five, seven and ten days of cultivation on PDA. **f,** The *VdAMP2* deletion mutant is not affected in microsclerotia formation. After ten days, colonies as shown in **e** were excised from plate and the tissue was ground to determine the number of microsclerotia per cm^2­^ using a haemocytometer.

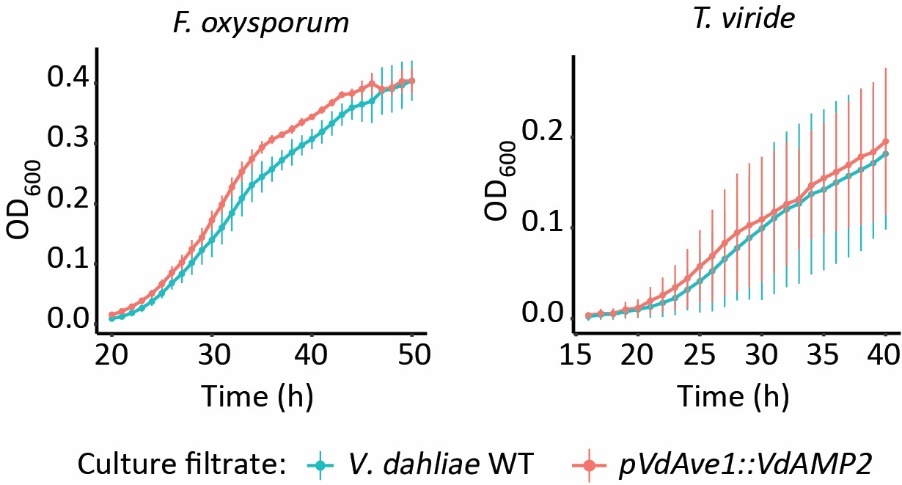

**Extended data Fig. 9: VdAMP2 does not inhibit fungal growth.** Growth of *F. oxysporum* and *T. viride* in filter-sterilized culture filtrates from *in vitro* grown wild-type *V. dahliae* and the *VdAMP2* expression transformant does not reveal antifungal activity of VdAMP2.

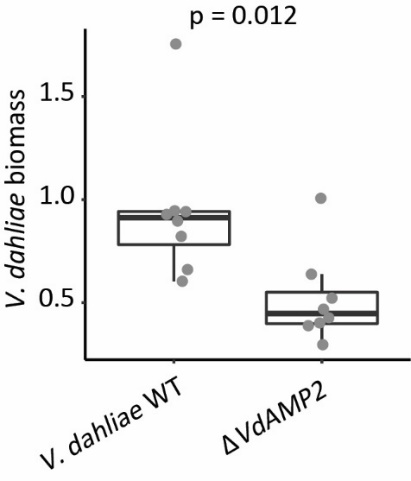

**Extended data Fig. 10: VdAMP2 contributes to host colonization.** Biomass of wild-type *V. dahliae* and the *VdAMP2* deletion mutant (Δ*VdAMP2*) in *A. thaliana* plants determined with real-time PCR at 27 days after sowing the seeds on soil containing microsclerotia of *V. dahliae* (unpaired student’s t-test; N=8).

**Extended data Table 1: DESeq2 output of differential abundance analysis of bacterial orders in tomato root microbiomes colonized by wild-type *V. dahliae* and the *VdAve1* deletion mutant**

| **Base mean** | **log2FC** | **lfcSE** | **stat** | **p value** | **Order** |
| --- | --- | --- | --- | --- | --- |
| 340.07 | 0.75 | 0.21 | 3.56 | 3.67E-04 | Bdellovibrionales |
| 265.53 | -0.60 | 0.19 | -3.24 | 1.18E-03 | Sphingomonadales |
| 143.41 | 0.97 | 0.31 | 3.17 | 1.55E-03 | Ktedonobacterales |
| 2126.65 | -0.32 | 0.12 | -2.57 | 1.02E-02 | Xanthomonadales |
| 146.92 | 1.30 | 0.58 | 2.25 | 2.42E-02 | Pseudomonadales |
| 705.03 | -0.32 | 0.15 | -2.17 | 3.01E-02 | Alphaproteobacteria incertae sedis |
| 104.59 | 0.70 | 0.34 | 2.06 | 3.97E-02 | Chlamydiales |
| 5.72 | -5.79 | 2.87 | -2.01 | 4.39E-02 | Acidimicrobiales |
| 28.94 | 1.95 | 0.98 | 2.00 | 4.54E-02 | Granulicella |
| 27.37 | -1.21 | 0.90 | -1.34 | 1.80E-01 | Hydrogenophilales |
| 444.45 | 0.24 | 0.18 | 1.34 | 1.80E-01 | Cytophagales |
| 25.12 | -1.00 | 0.74 | -1.34 | 1.81E-01 | Fimbriimonadales |
| 2.89 | -4.81 | 3.59 | -1.34 | 1.81E-01 | Sulfuricellales |
| 2.45 | 4.92 | 3.72 | 1.32 | 1.86E-01 | Methanosarcinales |
| 960.84 | 0.18 | 0.16 | 1.18 | 2.37E-01 | Actinomycetales |
| 308.73 | -0.21 | 0.19 | -1.10 | 2.72E-01 | Caulobacterales |
| 150.02 | -0.36 | 0.33 | -1.09 | 2.76E-01 | Spirochaetales |
| 442.03 | 0.17 | 0.17 | 1.04 | 3.00E-01 | Myxococcales |
| 1.74 | -4.07 | 3.93 | -1.04 | 3.00E-01 | Aeromonadales |
| 574.02 | -0.19 | 0.18 | -1.03 | 3.01E-01 | Flavobacteriales |
| 2112.11 | -0.09 | 0.09 | -1.03 | 3.04E-01 | Burkholderiales |
| 1.65 | -3.99 | 3.93 | -1.02 | 3.10E-01 | Thermoanaerobacterales |
| 1.48 | -3.84 | 3.93 | -0.98 | 3.29E-01 | Gammaproteobacteria incertae sedis |
| 1.14 | 3.82 | 3.95 | 0.97 | 3.34E-01 | Desulfuromonadales |
| 1.09 | 3.75 | 3.95 | 0.95 | 3.43E-01 | Legionellales |
| 37.82 | 0.50 | 0.55 | 0.91 | 3.65E-01 | Gaiellales |
| 301.06 | -0.17 | 0.20 | -0.85 | 3.97E-01 | Rhodospirillales |
| 160.75 | -0.20 | 0.24 | -0.81 | 4.17E-01 | Gp3 |
| 27.70 | -0.62 | 0.79 | -0.79 | 4.27E-01 | Bacteroidales |
| 19.95 | -0.68 | 0.86 | -0.79 | 4.32E-01 | Clostridiales |
| 2559.50 | -0.08 | 0.10 | -0.77 | 4.39E-01 | Sphingobacteriales |
| 1322.35 | 0.09 | 0.12 | 0.76 | 4.45E-01 | Rhizobiales |
| 14.69 | -2.52 | 3.33 | -0.76 | 4.49E-01 | Acidipila |
| 990.27 | -0.08 | 0.12 | -0.68 | 4.99E-01 | Telmatobacter |
| 6.39 | -1.42 | 2.24 | -0.63 | 5.26E-01 | Enterobacteriales |
| 23.72 | 0.50 | 0.80 | 0.62 | 5.35E-01 | Holophagales |
| 0.44 | 2.41 | 4.02 | 0.60 | 5.49E-01 | Sneathiellales |
| 3.16 | -1.93 | 3.23 | -0.60 | 5.51E-01 | Nitrososphaeraceae |
| 9.56 | -1.29 | 2.19 | -0.59 | 5.57E-01 | Candidatus_Koribacter |
| 270.02 | 0.12 | 0.21 | 0.56 | 5.76E-01 | Verrucomicrobiales |
| 4.54 | -1.57 | 2.87 | -0.55 | 5.85E-01 | Tepidisphaerales |
| 120.32 | 0.16 | 0.34 | 0.47 | 6.39E-01 | Terriglobus |
| 76.20 | -0.16 | 0.41 | -0.39 | 6.94E-01 | Solirubrobacterales |
| 66.75 | 0.14 | 0.37 | 0.37 | 7.10E-01 | Gallionellales |
| 53.14 | -0.55 | 1.49 | -0.37 | 7.12E-01 | Acidobacterium |
| 2.86 | 1.31 | 3.68 | 0.36 | 7.22E-01 | Gp2 |
| 37.32 | -0.15 | 0.52 | -0.30 | 7.68E-01 | Armatimonadales |
| 2.07 | 0.88 | 3.51 | 0.25 | 8.03E-01 | Lactobacillales |
| 11.06 | -0.45 | 1.82 | -0.25 | 8.05E-01 | Gemmatimonadales |
| 1.79 | 0.80 | 3.83 | 0.21 | 8.34E-01 | Anaerolineales |
| 372.06 | -0.04 | 0.18 | -0.21 | 8.36E-01 | Planctomycetales |
| 1.93 | 0.54 | 3.82 | 0.14 | 8.88E-01 | Chthonomonadales |
| 72.62 | -0.06 | 0.48 | -0.13 | 8.94E-01 | Opitutales |
| 14.47 | -0.19 | 1.60 | -0.12 | 9.07E-01 | Bacillales |
| 115.79 | -0.04 | 0.35 | -0.11 | 9.10E-01 | Rhodocyclales |
| 118.47 | 0.02 | 0.28 | 0.08 | 9.34E-01 | Gp1 |
| 72.58 | -0.02 | 0.37 | -0.06 | 9.56E-01 | Candidatus_Solibacter |
| 12.39 | 0.08 | 1.47 | 0.05 | 9.58E-01 | Nitrosomonadales |
| 11.43 | -0.05 | 1.27 | -0.04 | 9.69E-01 | Terrimicrobium |
| 224.78 | 0.01 | 0.76 | 0.01 | 9.94E-01 | Chromatiales |

**Extended data Table 2: DESeq2 output of differential abundance analysis of bacterial taxa (patristic distance<0.1) in cotton root microbiomes colonized by wild-type *V. dahliae* and the *VdAve1* deletion mutant**

| **Base mean** | **log2FC** | **lfcSE** | **stat** | **p value** | **Class** | **Order** | **Family** | **Genus** |
| --- | --- | --- | --- | --- | --- | --- | --- | --- |
| 131.29 | -2.30 | 0.39 | -5.82 | 5.98E-09 | Verrucomicrobiae | Verrucomicrobiales | Verrucomicrobiaceae | Luteolibacter |
| 64.48 | -3.60 | 0.86 | -4.18 | 2.93E-05 | Flavobacteriia | Flavobacteriales | Flavobacteriaceae | Chryseobacterium |
| 6.12 | 6.12 | 1.99 | 3.07 | 2.12E-03 | Alphaproteobacteria | Rhodospirillales | Rhodospirillaceae | NA |
| 144.44 | -1.24 | 0.41 | -3.01 | 2.63E-03 | Sphingobacteriia | Sphingobacteriales | Chitinophagaceae | Chitinophaga |
| 5.66 | -5.89 | 2.03 | -2.90 | 3.69E-03 | Alphaproteobacteria | Sphingomonadales | Sphingomonadaceae | NA |
| 5.47 | 5.97 | 2.07 | 2.88 | 3.95E-03 | Alphaproteobacteria | Rhodospirillales | Reyranella | NA |
| 1031.97 | -0.56 | 0.20 | -2.82 | 4.85E-03 | Betaproteobacteria | Burkholderiales | Oxalobacteraceae | NA |
| 4497.64 | 0.92 | 0.34 | 2.68 | 7.37E-03 | Gammaproteobacteria | Pseudomonadales | Pseudomonadaceae | Pseudomonas |
| 875.66 | 0.47 | 0.18 | 2.55 | 1.08E-02 | Sphingobacteriia | Sphingobacteriales | Sphingobacteriaceae | NA |
| 4.35 | -5.52 | 2.21 | -2.50 | 1.24E-02 | Deltaproteobacteria | Myxococcales | NA | NA |
| 37.69 | -2.74 | 1.09 | -2.50 | 1.25E-02 | Alphaproteobacteria | Rhizobiales | NA | NA |
| 5.77 | -5.92 | 2.66 | -2.23 | 2.60E-02 | Deltaproteobacteria | NA | NA | NA |
| 4.74 | 5.76 | 2.73 | 2.11 | 3.46E-02 | Alphaproteobacteria | Rhodospirillales | Rhodospirillaceae | NA |
| 149.00 | 1.06 | 0.52 | 2.06 | 3.99E-02 | Betaproteobacteria | Rhodocyclales | Rhodocyclaceae | NA |
| 4.13 | 5.56 | 2.82 | 1.97 | 4.86E-02 | Alphaproteobacteria | Alphaproteobacteria | Rhizomicrobium | NA |
| 6.60 | 6.24 | 3.17 | 1.97 | 4.93E-02 | Gammaproteobacteria | NA | NA | NA |
| 27.99 | -1.34 | 0.68 | -1.97 | 4.94E-02 | Sphingobacteriia | Sphingobacteriales | Chitinophagaceae | NA |
| 4.63 | 3.78 | 1.99 | 1.89 | 5.83E-02 | NA | NA | NA | NA |
| 3.84 | 5.46 | 2.88 | 1.89 | 5.84E-02 | Planctomycetia | Planctomycetales | Planctomycetaceae | NA |
| 3.79 | 5.44 | 2.90 | 1.88 | 6.04E-02 | Gammaproteobacteria | NA | NA | NA |
| 57.30 | 1.29 | 0.69 | 1.87 | 6.13E-02 | Flavobacteriia | Flavobacteriales | Flavobacteriaceae | Flavobacterium |
| 24.36 | -1.88 | 1.03 | -1.82 | 6.82E-02 | Actinobacteria | Actinomycetales | Actinospicaceae | Actinospica |
| 3.62 | 5.37 | 2.99 | 1.80 | 7.24E-02 | Armatimonadia | Armatimonadales | Armatimonadaceae | Armatimonas |
| 124.88 | 0.63 | 0.35 | 1.79 | 7.38E-02 | Armatimonadia | Armatimonadales | Armatimonadaceae | Armatimonas |
| 5.66 | -5.89 | 3.32 | -1.77 | 7.62E-02 | Sphingobacteriia | Sphingobacteriales | Chitinophagaceae | NA |
| 44.18 | 0.99 | 0.56 | 1.77 | 7.76E-02 | Betaproteobacteria | Neisseriales | Neisseriaceae | NA |
| 3.47 | 5.30 | 3.01 | 1.76 | 7.83E-02 | Cytophagia | Cytophagales | Cytophagaceae | NA |
| 5.48 | -5.85 | 3.35 | -1.75 | 8.08E-02 | Armatimonadia | Armatimonadales | Armatimonadaceae | Armatimonas |
| 21.72 | -1.36 | 0.79 | -1.74 | 8.26E-02 | Actinobacteria | Actinomycetales | NA | NA |
| 3.33 | 5.25 | 3.04 | 1.73 | 8.40E-02 | Planctomycetia | Planctomycetales | Planctomycetaceae | NA |
| 66.17 | -0.98 | 0.57 | -1.72 | 8.63E-02 | Alphaproteobacteria | Alphaproteobacteria | Rhizomicrobium | NA |
| 1715.94 | -0.68 | 0.40 | -1.67 | 9.41E-02 | Sphingobacteriia | Sphingobacteriales | Chitinophagaceae | Chitinophaga |
| 3.30 | -5.12 | 3.06 | -1.67 | 9.42E-02 | Alphaproteobacteria | NA | NA | NA |
| 26.54 | 1.25 | 0.75 | 1.66 | 9.69E-02 | NA | NA | NA | NA |
| 1402.13 | 0.63 | 0.38 | 1.65 | 9.95E-02 | Gammaproteobacteria | Enterobacteriales | Enterobacteriaceae | NA |
| 4.78 | -5.65 | 3.47 | -1.63 | 1.03E-01 | NA | NA | NA | NA |
| 2.93 | 5.07 | 3.13 | 1.62 | 1.05E-01 | Planctomycetia | Planctomycetales | Planctomycetaceae | Zavarzinella |
| 3.01 | 5.10 | 3.15 | 1.62 | 1.06E-01 | Deltaproteobacteria | Bdellovibrionales | Bdellovibrionaceae | Bdellovibrio |
| 2.80 | 5.00 | 3.17 | 1.58 | 1.15E-01 | Alphaproteobacteria | NA | NA | NA |
| 2.78 | 4.99 | 3.18 | 1.57 | 1.16E-01 | Alphaproteobacteria | Sphingomonadales | Sphingomonadaceae | NA |
| 57.03 | 0.85 | 0.55 | 1.55 | 1.21E-01 | Opitutae | Opitutales | Opitutaceae | NA |
| 3.91 | 5.48 | 3.63 | 1.51 | 1.31E-01 | Alphaproteobacteria | Rhodobacterales | Rhodobacteraceae | NA |
| 20.43 | -1.18 | 0.79 | -1.51 | 1.32E-01 | Flavobacteriia | Flavobacteriales | Cryomorphaceae | Fluviicola |
| 15.30 | -1.70 | 1.14 | -1.49 | 1.36E-01 | Alphaproteobacteria | Rhodospirillales | Rhodospirillaceae | NA |
| 13.18 | 2.40 | 1.62 | 1.48 | 1.39E-01 | Phycisphaerae | Tepidisphaerales | Tepidisphaeraceae | Tepidisphaera |
| 303.30 | 0.40 | 0.28 | 1.46 | 1.44E-01 | Gammaproteobacteria | Xanthomonadales | Xanthomonadaceae | NA |
| 109.68 | 0.52 | 0.36 | 1.44 | 1.51E-01 | Subdivision3 | NA | NA | NA |
| 56.39 | -0.98 | 0.69 | -1.43 | 1.54E-01 | Deltaproteobacteria | Bdellovibrionales | Bacteriovoracaceae | Peredibacter |
| 3.44 | 5.30 | 3.76 | 1.41 | 1.59E-01 | Acidobacteria_Gp1 | NA | NA | NA |
| 2.48 | -4.70 | 3.34 | -1.41 | 1.59E-01 | Sphingobacteriia | Sphingobacteriales | Chitinophagaceae | Flavisolibacter |
| 346.29 | -0.42 | 0.31 | -1.36 | 1.73E-01 | Gammaproteobacteria | NA | NA | NA |
| 100.78 | -0.55 | 0.40 | -1.36 | 1.75E-01 | Alphaproteobacteria | Sphingomonadales | Erythrobacteraceae | NA |
| 3.01 | 5.11 | 3.91 | 1.31 | 1.92E-01 | Alphaproteobacteria | Rhodospirillales | NA | NA |
| 3.18 | -5.06 | 3.88 | -1.31 | 1.92E-01 | Chlamydiia | Chlamydiales | Parachlamydiaceae | Neochlamydia |
| 2.04 | 4.55 | 3.53 | 1.29 | 1.97E-01 | Gammaproteobacteria | NA | NA | NA |
| 76.42 | 0.64 | 0.50 | 1.28 | 2.01E-01 | Alphaproteobacteria | Alphaproteobacteria | Rhizomicrobium | NA |
| 428.32 | 0.32 | 0.25 | 1.27 | 2.02E-01 | Betaproteobacteria | NA | NA | NA |
| 2.09 | -4.46 | 3.52 | -1.27 | 2.05E-01 | NA | NA | NA | NA |
| 2.73 | 4.96 | 3.92 | 1.26 | 2.06E-01 | NA | NA | NA | NA |
| 2.71 | 4.95 | 3.92 | 1.26 | 2.07E-01 | Actinobacteria | Solirubrobacterales | Conexibacteraceae | Conexibacter |
| 54.97 | -0.67 | 0.53 | -1.26 | 2.08E-01 | Sphingobacteriia | Sphingobacteriales | Chitinophagaceae | Arachidicoccus |
| 2.83 | -4.89 | 3.92 | -1.25 | 2.12E-01 | Actinobacteria | NA | NA | NA |
| 1.92 | 4.46 | 3.61 | 1.24 | 2.16E-01 | NA | NA | NA | NA |
| 2.44 | 4.80 | 3.93 | 1.22 | 2.21E-01 | Sphingobacteriia | Sphingobacteriales | Chitinophagaceae | Ferruginibacter |
| 155.60 | -0.39 | 0.33 | -1.20 | 2.30E-01 | Actinobacteria | Actinomycetales | Microbacteriaceae | NA |
| 2.48 | -4.70 | 3.92 | -1.20 | 2.31E-01 | NA | NA | NA | NA |
| 2.33 | -4.62 | 3.92 | -1.18 | 2.40E-01 | Betaproteobacteria | Burkholderiales | Burkholderiaceae | Cupriavidus |
| 2.13 | 4.59 | 3.93 | 1.17 | 2.43E-01 | Actinobacteria | Actinomycetales | Microbacteriaceae | NA |
| 2.11 | 4.59 | 3.93 | 1.17 | 2.43E-01 | Actinobacteria | Gaiellales | Gaiellaceae | Gaiella |
| 3.11 | -2.57 | 2.20 | -1.16 | 2.45E-01 | Flavobacteriia | Flavobacteriales | NA | NA |
| 9.79 | -1.98 | 1.70 | -1.16 | 2.45E-01 | Sphingobacteriia | Sphingobacteriales | Chitinophagaceae | Parafilimonas |
| 1.75 | 4.33 | 3.73 | 1.16 | 2.46E-01 | NA | NA | NA | NA |
| 2.20 | -4.53 | 3.93 | -1.15 | 2.48E-01 | Actinobacteria | Actinomycetales | NA | NA |
| 2.01 | 4.52 | 3.93 | 1.15 | 2.50E-01 | Opitutae | Opitutales | Opitutaceae | Opitutus |
| 1.81 | -4.25 | 3.70 | -1.15 | 2.50E-01 | Chlamydiia | Chlamydiales | Parachlamydiaceae | NA |
| 2378.54 | -0.20 | 0.18 | -1.13 | 2.59E-01 | Gammaproteobacteria | Xanthomonadales | Xanthomonadaceae | NA |
| 2.05 | -4.43 | 3.93 | -1.13 | 2.59E-01 | Alphaproteobacteria | Rhodospirillales | NA | NA |
| 2.05 | -4.43 | 3.93 | -1.13 | 2.59E-01 | Actinobacteria | Gaiellales | Gaiellaceae | Gaiella |
| 1.80 | 4.37 | 3.93 | 1.11 | 2.67E-01 | Alphaproteobacteria | Rhizobiales | Phyllobacteriaceae | NA |
| 1.66 | 4.24 | 3.82 | 1.11 | 2.68E-01 | NA | NA | NA | NA |
| 1.72 | 4.30 | 3.93 | 1.09 | 2.74E-01 | NA | NA | NA | NA |
| 1.70 | -4.16 | 3.81 | -1.09 | 2.75E-01 | Alphaproteobacteria | Rhizobiales | Hyphomicrobiaceae | Rhodomicrobium |
| 267.66 | -0.40 | 0.37 | -1.09 | 2.76E-01 | Alphaproteobacteria | Rhizobiales | NA | NA |
| 140.42 | -0.36 | 0.33 | -1.08 | 2.79E-01 | Actinobacteria | Actinomycetales | Mycobacteriaceae | Mycobacterium |
| 1.65 | 4.24 | 3.93 | 1.08 | 2.81E-01 | Deltaproteobacteria | Bdellovibrionales | Bdellovibrionaceae | Vampirovibrio |
| 60.52 | -0.73 | 0.68 | -1.07 | 2.82E-01 | Planctomycetia | Planctomycetales | Planctomycetaceae | NA |
| 1.66 | 4.22 | 3.94 | 1.07 | 2.83E-01 | Gammaproteobacteria | NA | NA | NA |
| 30.34 | 0.68 | 0.64 | 1.07 | 2.84E-01 | Alphaproteobacteria | NA | NA | NA |
| 26.47 | 0.72 | 0.67 | 1.07 | 2.85E-01 | NA | NA | NA | NA |
| 53.65 | 0.63 | 0.59 | 1.07 | 2.86E-01 | Betaproteobacteria | NA | NA | NA |
| 10.60 | 1.43 | 1.34 | 1.06 | 2.87E-01 | NA | NA | NA | NA |
| 1.58 | 4.18 | 3.94 | 1.06 | 2.89E-01 | Acidobacteria_Gp3 | NA | NA | NA |
| 1.58 | 4.18 | 3.94 | 1.06 | 2.89E-01 | Spartobacteria | NA | NA | NA |
| 1.58 | 4.18 | 3.94 | 1.06 | 2.89E-01 | NA | NA | NA | NA |
| 1.71 | -4.17 | 3.93 | -1.06 | 2.89E-01 | Acidobacteria_Gp2 | Gp2 | NA | NA |
| 1187.82 | 0.37 | 0.35 | 1.05 | 2.92E-01 | Alphaproteobacteria | Rhizobiales | NA | NA |
| 48.72 | 1.07 | 1.01 | 1.05 | 2.92E-01 | NA | NA | NA | NA |
| 9.61 | 1.58 | 1.51 | 1.05 | 2.94E-01 | Alphaproteobacteria | NA | NA | NA |
| 1.50 | 4.11 | 3.94 | 1.04 | 2.97E-01 | Subdivision3 | NA | NA | NA |
| 18.01 | 0.89 | 0.85 | 1.04 | 2.97E-01 | Flavobacteriia | Flavobacteriales | Cryomorphaceae | Fluviicola |
| 1.61 | -4.09 | 3.93 | -1.04 | 2.99E-01 | Subdivision3 | NA | NA | NA |
| 1.61 | -4.09 | 3.93 | -1.04 | 2.99E-01 | NA | NA | NA | NA |
| 72.22 | -0.47 | 0.45 | -1.03 | 3.01E-01 | Actinobacteria | Actinomycetales | Pseudonocardiaceae | Pseudonocardia |
| 1.59 | -4.06 | 3.93 | -1.03 | 3.02E-01 | Acidobacteria_Gp1 | Acidicapsa | NA | NA |
| 1.43 | 4.04 | 3.94 | 1.03 | 3.05E-01 | Alphaproteobacteria | NA | NA | NA |
| 1.43 | 4.04 | 3.94 | 1.03 | 3.05E-01 | Alphaproteobacteria | Rhodospirillales | Rhodospirillaceae | NA |
| 88.15 | -0.46 | 0.45 | -1.02 | 3.06E-01 | Gammaproteobacteria | Xanthomonadales | Sinobacteraceae | Steroidobacter |
| 585.84 | -0.21 | 0.21 | -1.01 | 3.12E-01 | Sphingobacteriia | Sphingobacteriales | Chitinophagaceae | NA |
| 75.67 | -0.45 | 0.45 | -1.01 | 3.14E-01 | Actinobacteria | Actinomycetales | Nocardioidaceae | Nocardioides |
| 14.04 | -1.26 | 1.27 | -1.00 | 3.20E-01 | Alphaproteobacteria | Rhodospirillales | Reyranella | NA |
| 65.55 | -0.67 | 0.68 | -0.99 | 3.22E-01 | Actinobacteria | Actinomycetales | NA | NA |
| 26.82 | 0.68 | 0.69 | 0.99 | 3.22E-01 | Spartobacteria | Terrimicrobium | NA | NA |
| 1.42 | -3.89 | 3.94 | -0.99 | 3.23E-01 | NA | NA | NA | NA |
| 1.29 | 3.89 | 3.94 | 0.99 | 3.24E-01 | Alphaproteobacteria | NA | NA | NA |
| 1.29 | 3.89 | 3.94 | 0.99 | 3.24E-01 | NA | NA | NA | NA |
| 1.29 | 3.89 | 3.94 | 0.99 | 3.24E-01 | Actinobacteria | Acidimicrobiales | Iamiaceae | Aquihabitans |
| 5.29 | 1.95 | 1.99 | 0.98 | 3.27E-01 | Alphaproteobacteria | Rhodospirillales | Rhodospirillaceae | NA |
| 21.64 | 1.20 | 1.23 | 0.97 | 3.30E-01 | Bacilli | Bacillales | Paenibacillaceae_1 | Paenibacillus |
| 58.31 | 0.64 | 0.66 | 0.97 | 3.31E-01 | Sphingobacteriia | Sphingobacteriales | Sphingobacteriaceae | Pedobacter |
| 28.48 | 0.70 | 0.72 | 0.97 | 3.33E-01 | Alphaproteobacteria | Rhodospirillales | Rhodospirillaceae | Dongia |
| 1.32 | -3.80 | 3.94 | -0.96 | 3.35E-01 | Actinobacteria | Actinomycetales | Thermomonosporaceae | Actinomadura |
| 1.32 | -3.80 | 3.94 | -0.96 | 3.35E-01 | Chlamydiia | Chlamydiales | Parachlamydiaceae | NA |
| 29.88 | -0.72 | 0.75 | -0.96 | 3.36E-01 | Actinobacteria | Actinomycetales | NA | NA |
| 1.20 | 3.79 | 3.95 | 0.96 | 3.37E-01 | Betaproteobacteria | NA | NA | NA |
| 1.20 | 3.79 | 3.95 | 0.96 | 3.37E-01 | Deltaproteobacteria | NA | NA | NA |
| 38.37 | -1.01 | 1.06 | -0.95 | 3.42E-01 | Deltaproteobacteria | Myxococcales | NA | NA |
| 13.89 | 1.98 | 2.08 | 0.95 | 3.42E-01 | Betaproteobacteria | NA | NA | NA |
| 1.18 | 3.73 | 3.95 | 0.95 | 3.44E-01 | Alphaproteobacteria | Rhizobiales | NA | NA |
| 45.77 | 0.55 | 0.59 | 0.94 | 3.46E-01 | Gammaproteobacteria | Xanthomonadales | Xanthomonadaceae | NA |
| 1.15 | 3.72 | 3.95 | 0.94 | 3.46E-01 | Acidobacteria_Gp2 | Gp2 | NA | NA |
| 1.15 | 3.72 | 3.95 | 0.94 | 3.46E-01 | Gemmatimonadetes | Gemmatimonadales | Gemmatimonadaceae | Gemmatimonas |
| 1.15 | 3.72 | 3.95 | 0.94 | 3.46E-01 | Sphingobacteriia | Sphingobacteriales | NA | NA |
| 1.15 | 3.72 | 3.95 | 0.94 | 3.46E-01 | NA | NA | NA | NA |
| 1.15 | 3.72 | 3.95 | 0.94 | 3.46E-01 | Acidobacteria_Gp3 | Candidatus_Solibacter | NA | NA |
| 5.98 | 1.80 | 1.92 | 0.94 | 3.49E-01 | Flavobacteriia | Flavobacteriales | Flavobacteriaceae | Flavobacterium |
| 15.88 | -1.15 | 1.23 | -0.94 | 3.49E-01 | NA | NA | NA | NA |
| 50.56 | -1.27 | 1.37 | -0.93 | 3.54E-01 | Cytophagia | Cytophagales | Cytophagaceae | Dyadobacter |
| 2058.72 | -0.34 | 0.37 | -0.92 | 3.55E-01 | Alphaproteobacteria | Rhizobiales | Rhizobiaceae | Rhizobium |
| 9.07 | -1.30 | 1.40 | -0.92 | 3.56E-01 | Gammaproteobacteria | NA | NA | NA |
| 1.05 | 3.59 | 3.95 | 0.91 | 3.63E-01 | Gammaproteobacteria | NA | NA | NA |
| 1229.31 | -0.42 | 0.47 | -0.90 | 3.69E-01 | Flavobacteriia | Flavobacteriales | Flavobacteriaceae | Flavobacterium |
| 1.00 | 3.53 | 3.95 | 0.89 | 3.72E-01 | Cytophagia | Cytophagales | Cytophagaceae | Adhaeribacter |
| 1.00 | 3.53 | 3.95 | 0.89 | 3.72E-01 | Ktedonobacteria | Thermogemmatisporales | Thermogemmatisporaceae | Thermogemmatispora |
| 1.09 | -3.52 | 3.95 | -0.89 | 3.73E-01 | Alphaproteobacteria | NA | NA | NA |
| 12.77 | 0.82 | 0.93 | 0.89 | 3.76E-01 | Gammaproteobacteria | NA | NA | NA |
| 14.33 | -1.06 | 1.21 | -0.88 | 3.78E-01 | Alphaproteobacteria | Rhodospirillales | Rhodospirillaceae | NA |
| 1.06 | -3.47 | 3.95 | -0.88 | 3.79E-01 | Clostridia | Thermoanaerobacterales | Thermoanaerobacteraceae | NA |
| 17.10 | 1.53 | 1.74 | 0.88 | 3.79E-01 | Acidobacteria_Gp1 | Gp1 | NA | NA |
| 327.11 | -0.33 | 0.38 | -0.88 | 3.82E-01 | Sphingobacteriia | Sphingobacteriales | Chitinophagaceae | Arachidicoccus |
| 1.02 | -3.44 | 3.95 | -0.87 | 3.84E-01 | Sphingobacteriia | Sphingobacteriales | Chitinophagaceae | NA |
| 93.73 | -0.36 | 0.41 | -0.87 | 3.85E-01 | Alphaproteobacteria | Rhizobiales | Xanthobacteraceae | Pseudolabrys |
| 9.50 | 1.05 | 1.21 | 0.86 | 3.87E-01 | Deltaproteobacteria | Bdellovibrionales | Bacteriovoracaceae | Bacteriovorax |
| 357.58 | -0.22 | 0.26 | -0.86 | 3.88E-01 | Actinobacteria | Actinomycetales | NA | NA |
| 0.95 | 3.41 | 3.96 | 0.86 | 3.89E-01 | Gammaproteobacteria | Xanthomonadales | Sinobacteraceae | Nevskia |
| 0.95 | 3.41 | 3.96 | 0.86 | 3.89E-01 | Verrucomicrobiae | Verrucomicrobiales | Verrucomicrobiaceae | NA |
| 0.95 | 3.41 | 3.96 | 0.86 | 3.89E-01 | Gemmatimonadetes | Gemmatimonadales | Gemmatimonadaceae | Gemmatimonas |
| 192.55 | -0.34 | 0.40 | -0.86 | 3.90E-01 | Deltaproteobacteria | Bdellovibrionales | Bdellovibrionaceae | Bdellovibrio |
| 31.15 | 0.56 | 0.65 | 0.86 | 3.91E-01 | Spartobacteria | NA | NA | NA |
| 828.22 | -0.28 | 0.33 | -0.86 | 3.92E-01 | Acidobacteria_Gp1 | NA | NA | NA |
| 11.29 | 0.83 | 0.97 | 0.86 | 3.92E-01 | Gammaproteobacteria | NA | NA | NA |
| 0.90 | 3.37 | 3.96 | 0.85 | 3.94E-01 | NA | NA | NA | NA |
| 4.81 | -2.09 | 2.45 | -0.85 | 3.95E-01 | Betaproteobacteria | NA | NA | NA |
| 311.37 | 0.24 | 0.28 | 0.85 | 3.95E-01 | Alphaproteobacteria | Alphaproteobacteria | Rhizomicrobium | NA |
| 11.56 | -1.55 | 1.83 | -0.85 | 3.96E-01 | Acidobacteria_Gp1 | Granulicella | NA | NA |
| 696.85 | 0.24 | 0.28 | 0.84 | 4.03E-01 | Gammaproteobacteria | Pseudomonadales | Pseudomonadaceae | Cellvibrio |
| 0.86 | 3.31 | 3.96 | 0.84 | 4.03E-01 | Betaproteobacteria | NA | NA | NA |
| 0.86 | 3.31 | 3.96 | 0.84 | 4.03E-01 | Planctomycetia | Planctomycetales | Planctomycetaceae | NA |
| 0.93 | -3.30 | 3.95 | -0.83 | 4.04E-01 | Alphaproteobacteria | Rhodospirillales | Acetobacteraceae | NA |
| 1281.84 | 0.16 | 0.20 | 0.83 | 4.09E-01 | NA | NA | NA | NA |
| 2.63 | 2.31 | 2.83 | 0.82 | 4.14E-01 | Chlamydiia | Chlamydiales | Parachlamydiaceae | Neochlamydia |
| 23.75 | -0.54 | 0.66 | -0.81 | 4.16E-01 | Acidobacteria_Gp3 | Gp3 | NA | NA |
| 41.17 | 0.51 | 0.63 | 0.81 | 4.17E-01 | NA | NA | NA | NA |
| 0.88 | -3.21 | 3.95 | -0.81 | 4.17E-01 | Gammaproteobacteria | NA | NA | NA |
| 0.88 | -3.21 | 3.95 | -0.81 | 4.17E-01 | Deltaproteobacteria | Myxococcales | Vulgatibacteraceae | Vulgatibacter |
| 0.88 | -3.21 | 3.95 | -0.81 | 4.17E-01 | NA | NA | NA | NA |
| 29.56 | 0.51 | 0.64 | 0.80 | 4.23E-01 | Planctomycetia | Planctomycetales | Planctomycetaceae | NA |
| 3.71 | -1.68 | 2.11 | -0.80 | 4.26E-01 | Alphaproteobacteria | NA | NA | NA |
| 6.24 | 1.51 | 1.90 | 0.79 | 4.27E-01 | Planctomycetia | Planctomycetales | Planctomycetaceae | Gimesia |
| 0.84 | -3.14 | 3.96 | -0.79 | 4.27E-01 | NA | NA | NA | NA |
| 250.12 | 0.27 | 0.34 | 0.79 | 4.27E-01 | Alphaproteobacteria | Sphingomonadales | Sphingomonadaceae | NA |
| 9.08 | 1.43 | 1.80 | 0.79 | 4.27E-01 | NA | NA | NA | NA |
| 0.75 | 3.11 | 3.97 | 0.78 | 4.33E-01 | Ktedonobacteria | Ktedonobacterales | Ktedonobacteraceae | Ktedonobacter |
| 0.75 | 3.11 | 3.97 | 0.78 | 4.33E-01 | NA | NA | NA | NA |
| 10.53 | 0.75 | 0.96 | 0.78 | 4.34E-01 | Verrucomicrobiae | Verrucomicrobiales | Verrucomicrobiaceae | Roseimicrobium |
| 0.79 | -3.06 | 3.96 | -0.77 | 4.40E-01 | Alphaproteobacteria | Sneathiellales | Sneathiellaceae | Taonella |
| 0.72 | 3.05 | 3.97 | 0.77 | 4.43E-01 | Armatimonadia | Armatimonadales | Armatimonadaceae | Armatimonas |
| 0.72 | 3.05 | 3.97 | 0.77 | 4.43E-01 | NA | NA | NA | NA |
| 0.72 | 3.05 | 3.97 | 0.77 | 4.43E-01 | NA | NA | NA | NA |
| 0.72 | 3.05 | 3.97 | 0.77 | 4.43E-01 | NA | NA | NA | NA |
| 0.72 | 3.05 | 3.97 | 0.77 | 4.43E-01 | Deltaproteobacteria | Myxococcales | NA | NA |
| 0.72 | 3.05 | 3.97 | 0.77 | 4.43E-01 | Alphaproteobacteria | Sneathiellales | Sneathiellaceae | Ferrovibrio |
| 0.78 | -3.03 | 3.96 | -0.77 | 4.44E-01 | Alphaproteobacteria | Rhodospirillales | Rhodospirillaceae | NA |
| 0.78 | -3.03 | 3.96 | -0.77 | 4.44E-01 | NA | NA | NA | NA |
| 7.35 | -2.14 | 2.82 | -0.76 | 4.48E-01 | Gammaproteobacteria | Pseudomonadales | Pseudomonadaceae | NA |
| 24.41 | 0.60 | 0.79 | 0.76 | 4.48E-01 | NA | NA | NA | NA |
| 0.71 | 2.98 | 3.97 | 0.75 | 4.53E-01 | Bacilli | Bacillales | NA | NA |
| 65.98 | 0.44 | 0.58 | 0.75 | 4.55E-01 | Spirochaetia | Spirochaetales | Spirochaetaceae | Spirochaeta |
| 12.10 | 1.01 | 1.36 | 0.75 | 4.56E-01 | Alphaproteobacteria | NA | NA | NA |
| 0.73 | -2.95 | 3.96 | -0.74 | 4.56E-01 | NA | NA | NA | NA |
| 0.73 | -2.95 | 3.96 | -0.74 | 4.56E-01 | NA | NA | NA | NA |
| 0.73 | -2.95 | 3.96 | -0.74 | 4.56E-01 | NA | NA | NA | NA |
| 34.56 | -0.48 | 0.64 | -0.74 | 4.59E-01 | Sphingobacteriia | Sphingobacteriales | Chitinophagaceae | NA |
| 36.23 | -0.47 | 0.63 | -0.74 | 4.61E-01 | NA | NA | NA | NA |
| 60.28 | -0.47 | 0.64 | -0.74 | 4.61E-01 | Actinobacteria | Solirubrobacterales | Conexibacteraceae | Conexibacter |
| 17.50 | -1.21 | 1.64 | -0.74 | 4.62E-01 | Actinobacteria | Actinomycetales | Nocardioidaceae | NA |
| 347.42 | 0.26 | 0.35 | 0.73 | 4.63E-01 | Flavobacteriia | Flavobacteriales | Flavobacteriaceae | Flavobacterium |
| 5.37 | -0.99 | 1.35 | -0.73 | 4.64E-01 | Bacilli | Lactobacillales | NA | NA |
| 108.17 | -0.29 | 0.40 | -0.73 | 4.64E-01 | Alphaproteobacteria | Sphingomonadales | Sphingomonadaceae | NA |
| 28.32 | -0.82 | 1.13 | -0.73 | 4.65E-01 | Acidobacteria_Gp1 | NA | NA | NA |
| 3.36 | 1.94 | 2.69 | 0.72 | 4.71E-01 | Subdivision3 | NA | NA | NA |
| 0.66 | -2.80 | 3.97 | -0.71 | 4.81E-01 | Gammaproteobacteria | NA | NA | NA |
| 0.60 | 2.79 | 3.98 | 0.70 | 4.83E-01 | Planctomycetia | Planctomycetales | Planctomycetaceae | Gemmata |
| 0.60 | 2.79 | 3.98 | 0.70 | 4.83E-01 | Planctomycetia | Planctomycetales | Planctomycetaceae | Zavarzinella |
| 162.57 | -0.39 | 0.55 | -0.70 | 4.85E-01 | Sphingobacteriia | Sphingobacteriales | Chitinophagaceae | NA |
| 5078.27 | 0.15 | 0.22 | 0.69 | 4.88E-01 | Subdivision3 | NA | NA | NA |
| 697.87 | -0.15 | 0.21 | -0.69 | 4.90E-01 | Sphingobacteriia | Sphingobacteriales | Sphingobacteriaceae | Mucilaginibacter |
| 0.57 | 2.73 | 3.99 | 0.69 | 4.93E-01 | Alphaproteobacteria | NA | NA | NA |
| 0.57 | 2.73 | 3.99 | 0.69 | 4.93E-01 | Opitutae | Puniceicoccales | Puniceicoccaceae | NA |
| 0.57 | 2.73 | 3.99 | 0.69 | 4.93E-01 | Planctomycetia | Planctomycetales | Planctomycetaceae | NA |
| 0.57 | 2.73 | 3.99 | 0.69 | 4.93E-01 | Actinobacteria | NA | NA | NA |
| 0.57 | 2.73 | 3.99 | 0.69 | 4.93E-01 | Deltaproteobacteria | Myxococcales | NA | NA |
| 0.57 | 2.73 | 3.99 | 0.69 | 4.93E-01 | Opitutae | NA | NA | NA |
| 19.88 | -0.88 | 1.29 | -0.68 | 4.94E-01 | Spirochaetia | Spirochaetales | Spirochaetaceae | Spirochaeta |
| 0.62 | -2.71 | 3.97 | -0.68 | 4.95E-01 | Alphaproteobacteria | NA | NA | NA |
| 0.62 | -2.71 | 3.97 | -0.68 | 4.95E-01 | Anaerolineae | Anaerolineales | Anaerolineaceae | NA |
| 591.95 | -0.16 | 0.25 | -0.67 | 5.04E-01 | Sphingobacteriia | Sphingobacteriales | Sphingobacteriaceae | NA |
| 19.26 | -0.51 | 0.77 | -0.66 | 5.07E-01 | Sphingobacteriia | Sphingobacteriales | Chitinophagaceae | NA |
| 0.59 | -2.63 | 3.98 | -0.66 | 5.08E-01 | NA | NA | NA | NA |
| 0.59 | -2.63 | 3.98 | -0.66 | 5.08E-01 | Actinobacteria | NA | NA | NA |
| 5.98 | 1.93 | 2.92 | 0.66 | 5.09E-01 | Cytophagia | Cytophagales | Cytophagaceae | Dyadobacter |
| 59.05 | 0.56 | 0.84 | 0.66 | 5.09E-01 | Verrucomicrobiae | Verrucomicrobiales | Verrucomicrobiaceae | Luteolibacter |
| 91.15 | -0.28 | 0.43 | -0.66 | 5.11E-01 | Gammaproteobacteria | Xanthomonadales | Xanthomonadaceae | NA |
| 160.43 | 0.31 | 0.49 | 0.64 | 5.19E-01 | Verrucomicrobiae | Verrucomicrobiales | Verrucomicrobiaceae | Luteolibacter |
| 4.89 | 1.58 | 2.47 | 0.64 | 5.22E-01 | NA | NA | NA | NA |
| 5.14 | -1.55 | 2.45 | -0.63 | 5.27E-01 | Gemmatimonadetes | Gemmatimonadales | Gemmatimonadaceae | Gemmatimonas |
| 0.53 | -2.47 | 3.99 | -0.62 | 5.36E-01 | NA | NA | NA | NA |
| 7.32 | 0.92 | 1.49 | 0.62 | 5.38E-01 | Actinobacteria | Actinomycetales | NA | NA |
| 58.17 | 0.36 | 0.59 | 0.60 | 5.46E-01 | Alphaproteobacteria | Rhizobiales | Rhodobiaceae | Parvibaculum |
| 0.45 | 2.38 | 4.01 | 0.59 | 5.52E-01 | NA | NA | NA | NA |
| 0.45 | 2.38 | 4.01 | 0.59 | 5.52E-01 | NA | NA | NA | NA |
| 0.45 | 2.38 | 4.01 | 0.59 | 5.52E-01 | Thermomicrobia | Sphaerobacterales | Sphaerobacteraceae | NA |
| 0.45 | 2.38 | 4.01 | 0.59 | 5.52E-01 | Alphaproteobacteria | NA | NA | NA |
| 0.45 | 2.38 | 4.01 | 0.59 | 5.52E-01 | NA | NA | NA | NA |
| 0.47 | 2.38 | 4.01 | 0.59 | 5.52E-01 | NA | NA | NA | NA |
| 16.66 | 0.76 | 1.29 | 0.59 | 5.54E-01 | Gemmatimonadetes | Gemmatimonadales | Gemmatimonadaceae | Gemmatimonas |
| 6.43 | -1.11 | 1.90 | -0.59 | 5.58E-01 | Cytophagia | Cytophagales | Flammeovirgaceae | NA |
| 140.47 | 0.20 | 0.34 | 0.58 | 5.59E-01 | Flavobacteriia | Flavobacteriales | Cryomorphaceae | Fluviicola |
| 0.43 | 2.32 | 4.01 | 0.58 | 5.63E-01 | Nitrososphaerales | Nitrososphaeraceae | Nitrososphaera | NA |
| 0.43 | 2.32 | 4.01 | 0.58 | 5.63E-01 | Nitrososphaerales | Nitrososphaeraceae | Nitrososphaera | NA |
| 0.43 | 2.32 | 4.01 | 0.58 | 5.63E-01 | Chlamydiia | Chlamydiales | NA | NA |
| 46.39 | 0.35 | 0.61 | 0.58 | 5.65E-01 | Acidobacteria_Gp1 | Gp1 | NA | NA |
| 0.47 | -2.30 | 4.00 | -0.57 | 5.66E-01 | Alphaproteobacteria | NA | NA | NA |
| 0.47 | -2.30 | 4.00 | -0.57 | 5.66E-01 | NA | NA | NA | NA |
| 0.47 | -2.30 | 4.00 | -0.57 | 5.66E-01 | NA | NA | NA | NA |
| 2.57 | 1.64 | 2.91 | 0.57 | 5.72E-01 | NA | NA | NA | NA |
| 41.57 | 0.38 | 0.68 | 0.56 | 5.75E-01 | Actinobacteria | Solirubrobacterales | NA | NA |
| 0.44 | -2.22 | 4.00 | -0.55 | 5.79E-01 | Gammaproteobacteria | NA | NA | NA |
| 0.44 | -2.22 | 4.00 | -0.55 | 5.79E-01 | Deltaproteobacteria | Myxococcales | NA | NA |
| 0.44 | -2.22 | 4.00 | -0.55 | 5.79E-01 | Caldilineae | Caldilineales | Caldilineaceae | Litorilinea |
| 0.44 | -2.22 | 4.00 | -0.55 | 5.79E-01 | Alphaproteobacteria | NA | NA | NA |
| 10.71 | 0.53 | 0.97 | 0.55 | 5.81E-01 | NA | NA | NA | NA |
| 12.00 | 0.51 | 0.92 | 0.55 | 5.82E-01 | Planctomycetia | Planctomycetales | Planctomycetaceae | NA |
| 26.94 | -0.65 | 1.20 | -0.54 | 5.87E-01 | Alphaproteobacteria | Rhodospirillales | Acetobacteraceae | Acidisoma |
| 6.93 | 1.28 | 2.36 | 0.54 | 5.89E-01 | Alphaproteobacteria | NA | NA | NA |
| 2.14 | 1.64 | 3.03 | 0.54 | 5.89E-01 | Gammaproteobacteria | NA | NA | NA |
| 60.85 | -0.29 | 0.53 | -0.54 | 5.89E-01 | Actinobacteria | Actinomycetales | Nocardiaceae | Nocardia |
| 25.73 | -0.44 | 0.82 | -0.54 | 5.91E-01 | NA | NA | NA | NA |
| 4.16 | -1.37 | 2.56 | -0.53 | 5.93E-01 | Gammaproteobacteria | NA | NA | NA |
| 5.43 | 1.29 | 2.44 | 0.53 | 5.98E-01 | Betaproteobacteria | NA | NA | NA |
| 397.91 | 0.15 | 0.29 | 0.53 | 5.98E-01 | Betaproteobacteria | NA | NA | NA |
| 337.26 | -0.16 | 0.30 | -0.53 | 5.99E-01 | Alphaproteobacteria | Sphingomonadales | NA | NA |
| 3.27 | 1.44 | 2.76 | 0.52 | 6.01E-01 | Armatimonadetes_gp5 | NA | NA | NA |
| 18.80 | 0.44 | 0.85 | 0.52 | 6.06E-01 | Sphingobacteriia | Sphingobacteriales | Chitinophagaceae | NA |
| 11.69 | 0.48 | 0.92 | 0.52 | 6.06E-01 | Alphaproteobacteria | Rhodospirillales | Rhodospirillaceae | NA |
| 30.19 | 0.32 | 0.63 | 0.51 | 6.10E-01 | Gammaproteobacteria | Pseudomonadales | Moraxellaceae | NA |
| 12.64 | 0.46 | 0.92 | 0.50 | 6.14E-01 | Planctomycetia | Planctomycetales | Planctomycetaceae | NA |
| 3.00 | 1.72 | 3.41 | 0.50 | 6.14E-01 | Chlamydiia | Chlamydiales | NA | NA |
| 759.19 | -0.10 | 0.20 | -0.50 | 6.15E-01 | Alphaproteobacteria | Rhizobiales | Bradyrhizobiaceae | NA |
| 1.33 | -1.77 | 3.53 | -0.50 | 6.17E-01 | Opitutae | Puniceicoccales | Puniceicoccaceae | NA |
| 6.09 | -0.97 | 1.94 | -0.50 | 6.18E-01 | NA | NA | NA | NA |
| 58.37 | 0.26 | 0.52 | 0.50 | 6.19E-01 | Opitutae | Opitutales | Opitutaceae | Opitutus |
| 15.69 | -0.47 | 0.95 | -0.49 | 6.23E-01 | Armatimonadia | Armatimonadales | Armatimonadaceae | Armatimonas |
| 16.08 | -0.38 | 0.79 | -0.49 | 6.27E-01 | Fimbriimonadia | Fimbriimonadales | Fimbriimonadaceae | Fimbriimonas |
| 22.93 | 0.80 | 1.65 | 0.48 | 6.28E-01 | Sphingobacteriia | Sphingobacteriales | Chitinophagaceae | Flavihumibacter |
| 2.64 | 1.38 | 2.85 | 0.48 | 6.28E-01 | Cytophagia | Cytophagales | Cytophagaceae | NA |
| 7.86 | 1.10 | 2.32 | 0.48 | 6.34E-01 | Gammaproteobacteria | NA | NA | NA |
| 0.35 | -1.88 | 4.03 | -0.47 | 6.41E-01 | Nitrososphaerales | Nitrososphaeraceae | Nitrososphaera | NA |
| 0.35 | -1.88 | 4.03 | -0.47 | 6.41E-01 | Alphaproteobacteria | NA | NA | NA |
| 0.35 | -1.88 | 4.03 | -0.47 | 6.41E-01 | NA | NA | NA | NA |
| 0.30 | 1.85 | 4.05 | 0.46 | 6.47E-01 | Verrucomicrobiae | Verrucomicrobiales | Verrucomicrobiaceae | Luteolibacter |
| 0.30 | 1.85 | 4.05 | 0.46 | 6.47E-01 | Acidobacteria_Gp1 | NA | NA | NA |
| 0.30 | 1.85 | 4.05 | 0.46 | 6.47E-01 | NA | NA | NA | NA |
| 4.33 | 1.43 | 3.13 | 0.46 | 6.48E-01 | NA | NA | NA | NA |
| 27.88 | 0.30 | 0.65 | 0.46 | 6.48E-01 | Subdivision3 | NA | NA | NA |
| 189.11 | -0.16 | 0.36 | -0.46 | 6.49E-01 | Alphaproteobacteria | Sphingomonadales | Sphingomonadaceae | Sphingobium |
| 4.85 | -1.13 | 2.51 | -0.45 | 6.51E-01 | Actinobacteria | Actinomycetales | Nocardiaceae | NA |
| 0.29 | 1.81 | 4.05 | 0.45 | 6.55E-01 | NA | NA | NA | NA |
| 0.29 | 1.81 | 4.05 | 0.45 | 6.55E-01 | Alphaproteobacteria | Caulobacterales | Caulobacteraceae | Asticcacaulis |
| 30.11 | 0.27 | 0.62 | 0.45 | 6.56E-01 | Actinobacteria | Gaiellales | Gaiellaceae | Gaiella |
| 477.48 | -0.15 | 0.35 | -0.44 | 6.57E-01 | Sphingobacteriia | Sphingobacteriales | Sphingobacteriaceae | NA |
| 9.35 | -0.63 | 1.43 | -0.44 | 6.57E-01 | Alphaproteobacteria | NA | NA | NA |
| 851.74 | -0.10 | 0.23 | -0.44 | 6.58E-01 | Sphingobacteriia | Sphingobacteriales | Chitinophagaceae | NA |
| 20.30 | 0.40 | 0.91 | 0.44 | 6.59E-01 | Planctomycetia | Planctomycetales | Planctomycetaceae | Schlesneria |
| 8.07 | 0.57 | 1.29 | 0.44 | 6.59E-01 | NA | NA | NA | NA |
| 2380.06 | 0.11 | 0.25 | 0.43 | 6.66E-01 | Alphaproteobacteria | Caulobacterales | Caulobacteraceae | Asticcacaulis |
| 23.64 | 0.36 | 0.85 | 0.43 | 6.70E-01 | Alphaproteobacteria | Rhizobiales | NA | NA |
| 251.96 | 0.14 | 0.32 | 0.42 | 6.71E-01 | Subdivision3 | NA | NA | NA |
| 0.31 | -1.71 | 4.04 | -0.42 | 6.72E-01 | NA | NA | NA | NA |
| 0.31 | -1.71 | 4.04 | -0.42 | 6.72E-01 | NA | NA | NA | NA |
| 1.24 | 1.65 | 3.92 | 0.42 | 6.74E-01 | NA | NA | NA | NA |
| 152.44 | -0.15 | 0.35 | -0.42 | 6.74E-01 | Alphaproteobacteria | Rhizobiales | Hyphomicrobiaceae | Hyphomicrobium |
| 116.14 | 0.19 | 0.46 | 0.42 | 6.77E-01 | Deltaproteobacteria | Myxococcales | Myxococcaceae | Aggregicoccus |
| 12.31 | -0.55 | 1.33 | -0.42 | 6.78E-01 | Actinobacteria | Solirubrobacterales | Conexibacteraceae | Conexibacter |
| 230.96 | 0.11 | 0.26 | 0.41 | 6.81E-01 | Alphaproteobacteria | Caulobacterales | Caulobacteraceae | NA |
| 81.95 | 0.16 | 0.40 | 0.41 | 6.82E-01 | Opitutae | Opitutales | Opitutaceae | Opitutus |
| 0.29 | -1.64 | 4.05 | -0.40 | 6.86E-01 | Planctomycetia | Planctomycetales | Planctomycetaceae | Rhodopirellula |
| 0.29 | -1.64 | 4.05 | -0.40 | 6.86E-01 | NA | NA | NA | NA |
| 0.29 | -1.64 | 4.05 | -0.40 | 6.86E-01 | Mollicutes | Entomoplasmatales | Entomoplasmataceae | Entomoplasma |
| 15.91 | -0.50 | 1.24 | -0.40 | 6.86E-01 | NA | NA | NA | NA |
| 2.79 | 1.13 | 2.83 | 0.40 | 6.89E-01 | Chthonomonadetes | Chthonomonadales | Chthonomonadaceae | Chthonomonas |
| 24.54 | -0.31 | 0.77 | -0.40 | 6.90E-01 | Cytophagia | Cytophagales | Cytophagaceae | Cytophaga |
| 13.77 | -0.35 | 0.89 | -0.39 | 6.95E-01 | NA | NA | NA | NA |
| 13.20 | -0.53 | 1.35 | -0.39 | 6.95E-01 | Sphingobacteriia | Sphingobacteriales | Chitinophagaceae | NA |
| 80.21 | 0.18 | 0.45 | 0.39 | 6.96E-01 | Opitutae | Opitutales | Opitutaceae | Opitutus |
| 385.64 | -0.10 | 0.26 | -0.39 | 6.99E-01 | Alphaproteobacteria | Rhodospirillales | Acetobacteraceae | NA |
| 20.13 | -0.28 | 0.73 | -0.38 | 7.01E-01 | Alphaproteobacteria | Rhodospirillales | NA | NA |
| 63.63 | -0.19 | 0.50 | -0.38 | 7.01E-01 | Alphaproteobacteria | Rhizobiales | NA | NA |
| 61.55 | -0.20 | 0.52 | -0.38 | 7.05E-01 | Sphingobacteriia | Sphingobacteriales | Chitinophagaceae | NA |
| 40.47 | 0.29 | 0.79 | 0.37 | 7.08E-01 | Chlamydiia | Chlamydiales | NA | NA |
| 12.45 | -0.47 | 1.27 | -0.37 | 7.11E-01 | Sphingobacteriia | Sphingobacteriales | Chitinophagaceae | Ferruginibacter |
| 9.41 | 0.84 | 2.31 | 0.36 | 7.15E-01 | Acidobacteria_Gp1 | Telmatobacter | NA | NA |
| 81.50 | 0.16 | 0.43 | 0.36 | 7.16E-01 | Alphaproteobacteria | Rhodospirillales | Rhodospirillaceae | NA |
| 80.30 | -0.20 | 0.54 | -0.36 | 7.16E-01 | NA | NA | NA | NA |
| 24.01 | 0.31 | 0.87 | 0.36 | 7.18E-01 | Betaproteobacteria | Neisseriales | Neisseriaceae | Snodgrassella |
| 5.18 | -0.89 | 2.48 | -0.36 | 7.20E-01 | Actinobacteria | Actinomycetales | Nocardiaceae | Rhodococcus |
| 39.96 | 0.25 | 0.71 | 0.36 | 7.21E-01 | NA | NA | NA | NA |
| 4.41 | 1.11 | 3.13 | 0.36 | 7.22E-01 | Deltaproteobacteria | Bdellovibrionales | Bdellovibrionaceae | Bdellovibrio |
| 105.93 | -0.14 | 0.40 | -0.34 | 7.32E-01 | NA | NA | NA | NA |
| 1063.04 | -0.10 | 0.29 | -0.34 | 7.32E-01 | Betaproteobacteria | NA | NA | NA |
| 2.25 | 1.03 | 3.03 | 0.34 | 7.33E-01 | NA | NA | NA | NA |
| 21.65 | -0.42 | 1.24 | -0.34 | 7.34E-01 | Deltaproteobacteria | Bdellovibrionales | Bdellovibrionaceae | Vampirovibrio |
| 830.09 | 0.09 | 0.25 | 0.34 | 7.36E-01 | Alphaproteobacteria | NA | NA | NA |
| 35.87 | 0.20 | 0.59 | 0.34 | 7.38E-01 | Actinobacteria | Actinomycetales | Nocardioidaceae | Nocardioides |
| 7.46 | 0.51 | 1.53 | 0.33 | 7.38E-01 | Cytophagia | Cytophagales | NA | NA |
| 135.59 | 0.14 | 0.42 | 0.33 | 7.41E-01 | Acidobacteria_Gp1 | NA | NA | NA |
| 6.43 | 0.79 | 2.41 | 0.33 | 7.42E-01 | Alphaproteobacteria | Caulobacterales | Caulobacteraceae | Brevundimonas |
| 3008.32 | 0.07 | 0.22 | 0.33 | 7.43E-01 | Betaproteobacteria | Burkholderiales | Burkholderiaceae | NA |
| 45.32 | -0.16 | 0.51 | -0.32 | 7.49E-01 | Sphingobacteriia | Sphingobacteriales | Chitinophagaceae | NA |
| 3.45 | 0.86 | 2.70 | 0.32 | 7.51E-01 | Armatimonadetes_gp4 | NA | NA | NA |
| 11.07 | -0.41 | 1.31 | -0.32 | 7.53E-01 | NA | NA | NA | NA |
| 53.40 | -0.18 | 0.59 | -0.31 | 7.57E-01 | Acidobacteria_Gp3 | NA | NA | NA |
| 10.99 | -0.43 | 1.39 | -0.31 | 7.58E-01 | Armatimonadia | Armatimonadales | Armatimonadaceae | Armatimonas |
| 1.52 | 1.18 | 3.86 | 0.31 | 7.60E-01 | Planctomycetia | Planctomycetales | Planctomycetaceae | Pirellula |
| 38.16 | -0.19 | 0.61 | -0.31 | 7.60E-01 | Actinobacteria | Actinomycetales | Thermomonosporaceae | NA |
| 8.53 | -0.36 | 1.18 | -0.31 | 7.60E-01 | Methanomicrobia | Methanosarcinales | NA | NA |
| 3.68 | 0.99 | 3.26 | 0.30 | 7.61E-01 | Subdivision3 | NA | NA | NA |
| 7.45 | 0.72 | 2.40 | 0.30 | 7.63E-01 | Sphingobacteriia | Sphingobacteriales | Chitinophagaceae | NA |
| 1.88 | 1.14 | 3.84 | 0.30 | 7.66E-01 | Chlamydiia | Chlamydiales | Simkaniaceae | Simkania |
| 10.56 | -0.40 | 1.35 | -0.30 | 7.66E-01 | NA | NA | NA | NA |
| 3.64 | 0.97 | 3.27 | 0.30 | 7.66E-01 | NA | NA | NA | NA |
| 1.68 | 1.14 | 3.85 | 0.30 | 7.68E-01 | Alphaproteobacteria | Rhodospirillales | NA | NA |
| 33.62 | 0.17 | 0.59 | 0.30 | 7.68E-01 | Planctomycetia | Planctomycetales | Planctomycetaceae | NA |
| 33.97 | 0.18 | 0.60 | 0.29 | 7.69E-01 | Gemmatimonadetes | Gemmatimonadales | Gemmatimonadaceae | Gemmatimonas |
| 9.38 | 0.43 | 1.47 | 0.29 | 7.70E-01 | Alphaproteobacteria | NA | NA | NA |
| 5.50 | -0.56 | 1.99 | -0.28 | 7.78E-01 | Alphaproteobacteria | Rhodospirillales | Rhodospirillaceae | NA |
| 7.27 | -0.53 | 1.90 | -0.28 | 7.78E-01 | Actinobacteria | Solirubrobacterales | Patulibacteraceae | Patulibacter |
| 1.81 | -1.08 | 3.84 | -0.28 | 7.78E-01 | Bacilli | Bacillales | NA | NA |
| 5.07 | -0.85 | 3.05 | -0.28 | 7.82E-01 | Acidobacteria_Gp1 | Gp1 | NA | NA |
| 256.89 | -0.09 | 0.33 | -0.27 | 7.89E-01 | Betaproteobacteria | Burkholderiales | Alcaligenaceae | NA |
| 1.38 | -1.03 | 3.88 | -0.26 | 7.91E-01 | Alphaproteobacteria | NA | NA | NA |
| 5.88 | -0.78 | 2.97 | -0.26 | 7.92E-01 | Subdivision3 | NA | NA | NA |
| 3.06 | -0.71 | 2.79 | -0.26 | 7.98E-01 | NA | NA | NA | NA |
| 8.14 | -0.35 | 1.40 | -0.25 | 8.01E-01 | NA | NA | NA | NA |
| 0.92 | -0.97 | 3.94 | -0.25 | 8.05E-01 | NA | NA | NA | NA |
| 0.92 | -0.97 | 3.94 | -0.25 | 8.05E-01 | Gammaproteobacteria | NA | NA | NA |
| 3.61 | 0.80 | 3.27 | 0.25 | 8.06E-01 | Acidobacteria_Gp6 | Gp6 | NA | NA |
| 1.61 | 0.81 | 3.31 | 0.25 | 8.06E-01 | Alphaproteobacteria | NA | NA | NA |
| 20.67 | 0.18 | 0.74 | 0.25 | 8.06E-01 | NA | NA | NA | NA |
| 7.42 | 0.46 | 1.88 | 0.24 | 8.07E-01 | Betaproteobacteria | NA | NA | NA |
| 3.70 | -0.64 | 2.66 | -0.24 | 8.08E-01 | NA | NA | NA | NA |
| 464.52 | -0.05 | 0.22 | -0.24 | 8.09E-01 | Alphaproteobacteria | Rhizobiales | NA | NA |
| 5.57 | 0.31 | 1.28 | 0.24 | 8.10E-01 | Actinobacteria | Solirubrobacterales | Solirubrobacteraceae | Solirubrobacter |
| 11.59 | 0.22 | 0.93 | 0.24 | 8.14E-01 | Alphaproteobacteria | Rhodospirillales | Rhodospirillaceae | NA |
| 1.77 | 0.89 | 3.84 | 0.23 | 8.16E-01 | Sphingobacteriia | Sphingobacteriales | Chitinophagaceae | NA |
| 160.15 | 0.11 | 0.48 | 0.22 | 8.22E-01 | Sphingobacteriia | Sphingobacteriales | Chitinophagaceae | Taibaiella |
| 52.34 | -0.15 | 0.70 | -0.22 | 8.27E-01 | Alphaproteobacteria | Caulobacterales | Caulobacteraceae | Phenylobacterium |
| 2.38 | 0.78 | 3.62 | 0.21 | 8.30E-01 | Actinobacteria | Acidimicrobiales | NA | NA |
| 8.98 | -0.31 | 1.46 | -0.21 | 8.30E-01 | Gemmatimonadetes | Gemmatimonadales | Gemmatimonadaceae | Gemmatimonas |
| 34.93 | -0.14 | 0.64 | -0.21 | 8.31E-01 | Gammaproteobacteria | Xanthomonadales | Sinobacteraceae | NA |
| 3.25 | -0.58 | 2.74 | -0.21 | 8.31E-01 | Clostridia | NA | NA | NA |
| 62.90 | -0.11 | 0.50 | -0.21 | 8.33E-01 | NA | NA | NA | NA |
| 10.85 | 0.39 | 1.85 | 0.21 | 8.35E-01 | Planctomycetia | Planctomycetales | Planctomycetaceae | Planctopirus |
| 9.02 | 0.38 | 1.89 | 0.20 | 8.42E-01 | Planctomycetia | Planctomycetales | Planctomycetaceae | Gemmata |
| 2.07 | 0.74 | 3.76 | 0.20 | 8.44E-01 | Spartobacteria | Terrimicrobium | NA | NA |
| 1.23 | -0.76 | 3.89 | -0.20 | 8.44E-01 | Actinobacteria | Acidimicrobiales | NA | NA |
| 62.24 | 0.09 | 0.47 | 0.19 | 8.49E-01 | Alphaproteobacteria | Rhodospirillales | Acetobacteraceae | NA |
| 34.08 | -0.14 | 0.72 | -0.19 | 8.49E-01 | Alphaproteobacteria | Sphingomonadales | NA | NA |
| 2.61 | -0.67 | 3.54 | -0.19 | 8.50E-01 | Deltaproteobacteria | Myxococcales | Labilitrichaceae | Labilithrix |
| 109.21 | 0.08 | 0.40 | 0.19 | 8.51E-01 | Alphaproteobacteria | Rhizobiales | Roseiarcaceae | Roseiarcus |
| 18.33 | -0.14 | 0.75 | -0.18 | 8.54E-01 | Alphaproteobacteria | Sphingomonadales | Sphingomonadaceae | NA |
| 26.92 | 0.12 | 0.63 | 0.18 | 8.55E-01 | Planctomycetia | Planctomycetales | Planctomycetaceae | NA |
| 3.11 | 0.59 | 3.39 | 0.17 | 8.61E-01 | Actinobacteria | Solirubrobacterales | NA | NA |
| 78.41 | 0.09 | 0.51 | 0.17 | 8.66E-01 | Alphaproteobacteria | Sphingomonadales | Sphingomonadaceae | NA |
| 5.02 | -0.34 | 2.05 | -0.17 | 8.68E-01 | Planctomycetia | Planctomycetales | Planctomycetaceae | NA |
| 33.33 | 0.10 | 0.62 | 0.16 | 8.70E-01 | Sphingobacteriia | Sphingobacteriales | Chitinophagaceae | NA |
| 609.60 | 0.04 | 0.28 | 0.16 | 8.76E-01 | Cytophagia | Cytophagales | Cytophagaceae | Cytophaga |
| 5.36 | 0.30 | 1.98 | 0.15 | 8.81E-01 | Alphaproteobacteria | Rhodobacterales | Rhodobacteraceae | NA |
| 44.88 | -0.09 | 0.62 | -0.15 | 8.84E-01 | Alphaproteobacteria | Caulobacterales | Caulobacteraceae | NA |
| 16.70 | 0.18 | 1.24 | 0.15 | 8.84E-01 | Gemmatimonadetes | Gemmatimonadales | Gemmatimonadaceae | Gemmatimonas |
| 7.22 | 0.27 | 1.91 | 0.14 | 8.87E-01 | NA | NA | NA | NA |
| 5.38 | 0.41 | 3.06 | 0.14 | 8.93E-01 | Betaproteobacteria | Burkholderiales | NA | NA |
| 3.64 | 0.36 | 2.73 | 0.13 | 8.95E-01 | Spirochaetia | Spirochaetales | Spirochaetaceae | Spirochaeta |
| 1.81 | 0.42 | 3.21 | 0.13 | 8.95E-01 | Alphaproteobacteria | NA | NA | NA |
| 25.20 | 0.11 | 0.83 | 0.13 | 8.95E-01 | Acidobacteria_Gp1 | NA | NA | NA |
| 54.00 | 0.06 | 0.48 | 0.13 | 8.99E-01 | Holophagae | Holophagales | Holophagaceae | Geothrix |
| 89.49 | -0.05 | 0.41 | -0.13 | 8.99E-01 | NA | NA | NA | NA |
| 6.68 | 0.30 | 2.39 | 0.13 | 8.99E-01 | Sphingobacteriia | Sphingobacteriales | Chitinophagaceae | Taibaiella |
| 20.05 | 0.11 | 0.86 | 0.13 | 9.00E-01 | Planctomycetia | Planctomycetales | Planctomycetaceae | NA |
| 154.74 | -0.04 | 0.34 | -0.12 | 9.02E-01 | Verrucomicrobiae | Verrucomicrobiales | Verrucomicrobiaceae | Prosthecobacter |
| 5.50 | 0.20 | 1.61 | 0.12 | 9.02E-01 | NA | NA | NA | NA |
| 17.52 | -0.10 | 0.82 | -0.12 | 9.04E-01 | NA | NA | NA | NA |
| 808.18 | -0.03 | 0.25 | -0.12 | 9.06E-01 | Alphaproteobacteria | Rhizobiales | Hyphomicrobiaceae | Devosia |
| 129.47 | 0.06 | 0.49 | 0.12 | 9.06E-01 | Planctomycetia | Planctomycetales | Planctomycetaceae | NA |
| 52.98 | -0.08 | 0.72 | -0.12 | 9.06E-01 | NA | NA | NA | NA |
| 8.94 | 0.27 | 2.29 | 0.12 | 9.07E-01 | Acidobacteria_Gp1 | NA | NA | NA |
| 245.12 | -0.05 | 0.42 | -0.11 | 9.10E-01 | Spartobacteria | NA | NA | NA |
| 2.45 | -0.40 | 3.59 | -0.11 | 9.11E-01 | Gammaproteobacteria | Xanthomonadales | Xanthomonadaceae | NA |
| 1.02 | -0.42 | 3.91 | -0.11 | 9.14E-01 | Subdivision3 | NA | NA | NA |
| 14.46 | 0.09 | 0.88 | 0.10 | 9.17E-01 | NA | NA | NA | NA |
| 12.99 | -0.18 | 1.77 | -0.10 | 9.18E-01 | Chlamydiia | Chlamydiales | NA | NA |
| 42.05 | 0.07 | 0.69 | 0.10 | 9.18E-01 | Alphaproteobacteria | NA | NA | NA |
| 265.72 | 0.05 | 0.49 | 0.10 | 9.19E-01 | Flavobacteriia | Flavobacteriales | Flavobacteriaceae | Flavobacterium |
| 10.27 | 0.10 | 1.01 | 0.10 | 9.20E-01 | Deltaproteobacteria | Bdellovibrionales | Bdellovibrionaceae | Bdellovibrio |
| 23.32 | 0.07 | 0.71 | 0.10 | 9.21E-01 | Verrucomicrobiae | Verrucomicrobiales | Verrucomicrobiaceae | NA |
| 5.56 | 0.30 | 3.03 | 0.10 | 9.22E-01 | Alphaproteobacteria | Caulobacterales | Caulobacteraceae | Phenylobacterium |
| 27.33 | -0.07 | 0.69 | -0.09 | 9.25E-01 | Betaproteobacteria | NA | NA | NA |
| 37.12 | -0.06 | 0.65 | -0.09 | 9.29E-01 | Acidobacteria_Gp3 | Gp3 | NA | NA |
| 5.69 | -0.26 | 2.99 | -0.09 | 9.31E-01 | Sphingobacteriia | Sphingobacteriales | Chitinophagaceae | NA |
| 1.35 | -0.30 | 3.52 | -0.09 | 9.31E-01 | Alphaproteobacteria | NA | NA | NA |
| 6.52 | -0.16 | 1.92 | -0.09 | 9.32E-01 | Deltaproteobacteria | Bdellovibrionales | Bdellovibrionaceae | Bdellovibrio |
| 451.20 | 0.04 | 0.42 | 0.08 | 9.33E-01 | Alphaproteobacteria | Caulobacterales | Caulobacteraceae | Caulobacter |
| 4.18 | 0.17 | 2.11 | 0.08 | 9.34E-01 | Gammaproteobacteria | NA | NA | NA |
| 1.04 | 0.32 | 3.90 | 0.08 | 9.35E-01 | Alphaproteobacteria | NA | NA | NA |
| 9.32 | -0.14 | 1.80 | -0.08 | 9.38E-01 | Acidobacteria_Gp2 | Gp2 | NA | NA |
| 1.68 | -0.30 | 3.84 | -0.08 | 9.38E-01 | Betaproteobacteria | Rhodocyclales | Rhodocyclaceae | NA |
| 19.09 | 0.08 | 0.98 | 0.08 | 9.39E-01 | Spirochaetia | Spirochaetales | Spirochaetaceae | NA |
| 5.62 | 0.13 | 1.65 | 0.08 | 9.39E-01 | Planctomycetia | Planctomycetales | Planctomycetaceae | Zavarzinella |
| 42.97 | -0.04 | 0.53 | -0.07 | 9.41E-01 | Betaproteobacteria | Burkholderiales | Alcaligenaceae | NA |
| 3.72 | 0.24 | 3.26 | 0.07 | 9.41E-01 | Alphaproteobacteria | Rhodospirillales | Rhodospirillaceae | NA |
| 1.92 | -0.28 | 3.82 | -0.07 | 9.42E-01 | Sphingobacteriia | Sphingobacteriales | Chitinophagaceae | NA |
| 35.79 | -0.06 | 0.76 | -0.07 | 9.42E-01 | Alphaproteobacteria | Sphingomonadales | NA | NA |
| 69.16 | -0.03 | 0.49 | -0.07 | 9.44E-01 | Alphaproteobacteria | Rhizobiales | NA | NA |
| 1.94 | 0.27 | 3.82 | 0.07 | 9.44E-01 | Actinobacteria | Acidimicrobiales | Iamiaceae | NA |
| 7.67 | 0.10 | 1.47 | 0.07 | 9.46E-01 | Gammaproteobacteria | Oceanospirillales | Oceanospirillaceae | NA |
| 32.37 | 0.04 | 0.65 | 0.07 | 9.47E-01 | Gammaproteobacteria | Xanthomonadales | Sinobacteraceae | Solimonas |
| 2.23 | 0.24 | 3.69 | 0.06 | 9.49E-01 | NA | NA | NA | NA |
| 9.78 | -0.11 | 1.85 | -0.06 | 9.52E-01 | Actinobacteria | Actinomycetales | NA | NA |
| 3.12 | -0.13 | 2.29 | -0.06 | 9.54E-01 | NA | NA | NA | NA |
| 1.96 | -0.22 | 3.82 | -0.06 | 9.55E-01 | Actinobacteria | Gaiellales | Gaiellaceae | Gaiella |
| 35.51 | -0.04 | 0.67 | -0.05 | 9.58E-01 | Planctomycetia | Planctomycetales | Planctomycetaceae | NA |
| 3.34 | -0.14 | 2.75 | -0.05 | 9.59E-01 | Alphaproteobacteria | NA | NA | NA |
| 3.41 | 0.17 | 3.32 | 0.05 | 9.60E-01 | NA | NA | NA | NA |
| 99.54 | -0.02 | 0.43 | -0.05 | 9.61E-01 | Alphaproteobacteria | Rhodospirillales | Rhodospirillaceae | Dongia |
| 4.42 | 0.12 | 2.60 | 0.05 | 9.62E-01 | Alphaproteobacteria | Rhodospirillales | NA | NA |
| 3.35 | -0.15 | 3.35 | -0.04 | 9.65E-01 | Acidobacteria_Gp1 | NA | NA | NA |
| 12.20 | 0.06 | 1.39 | 0.04 | 9.66E-01 | Chlamydiia | Chlamydiales | Parachlamydiaceae | Parachlamydia |
| 6.54 | 0.09 | 2.41 | 0.04 | 9.70E-01 | Actinobacteria | Acidimicrobiales | NA | NA |
| 28.73 | 0.03 | 0.75 | 0.04 | 9.71E-01 | NA | NA | NA | NA |
| 41.40 | 0.02 | 0.65 | 0.03 | 9.75E-01 | Subdivision3 | NA | NA | NA |
| 3.02 | -0.07 | 2.31 | -0.03 | 9.76E-01 | NA | NA | NA | NA |
| 14.04 | -0.04 | 1.25 | -0.03 | 9.76E-01 | Acidobacteria_Gp3 | Gp3 | NA | NA |
| 22.45 | 0.02 | 0.69 | 0.03 | 9.77E-01 | Deltaproteobacteria | Bdellovibrionales | Bdellovibrionaceae | Bdellovibrio |
| 348.50 | -0.01 | 0.37 | -0.03 | 9.78E-01 | Opitutae | Opitutales | Opitutaceae | Opitutus |
| 405.64 | -0.01 | 0.33 | -0.03 | 9.78E-01 | NA | NA | NA | NA |
| 48.01 | -0.01 | 0.57 | -0.02 | 9.80E-01 | Sphingobacteriia | Sphingobacteriales | Chitinophagaceae | NA |
| 23.89 | -0.03 | 1.15 | -0.02 | 9.80E-01 | Actinobacteria | Actinomycetales | Micrococcaceae | NA |
| 3.82 | 0.08 | 3.24 | 0.02 | 9.81E-01 | Alphaproteobacteria | Rhodospirillales | Rhodospirillaceae | NA |
| 19.28 | -0.02 | 0.83 | -0.02 | 9.82E-01 | Alphaproteobacteria | Caulobacterales | Caulobacteraceae | Phenylobacterium |
| 5.77 | 0.02 | 1.60 | 0.01 | 9.90E-01 | Alphaproteobacteria | NA | NA | NA |
| 0.58 | 0.05 | 4.01 | 0.01 | 9.90E-01 | Acidobacteria_Gp14 | Gp14 | NA | NA |
| 87.39 | 0.00 | 0.39 | -0.01 | 9.91E-01 | Sphingobacteriia | Sphingobacteriales | Chitinophagaceae | Sediminibacterium |
| 12.57 | -0.01 | 1.39 | -0.01 | 9.92E-01 | NA | NA | NA | NA |
| 5.79 | -0.01 | 1.64 | -0.01 | 9.93E-01 | NA | NA | NA | NA |
| 8.90 | 0.01 | 1.41 | 0.01 | 9.94E-01 | NA | NA | NA | NA |
| 36.54 | 0.00 | 0.90 | 0.00 | 1.00E+00 | NA | NA | NA | NA |

**Extended data Table 3: Differential abundance analysis of bacterial orders following combination of tomato and cotton samples based on infection by wild-type *V. dahliae* and the *VdAve1* deletion mutant**

| **Average relative abundance (%)** | **log2FC** | **p value** | **Order** |
| --- | --- | --- | --- |
| 1.84 | -0.37 | 3.41E-03 | Sphingomonadales |
| 0.08 | -0.71 | 1.76E-02 | Fimbriimonadales |
| 5.30 | 1.05 | 2.35E-02 | Pseudomonadales |
| 13.61 | -0.18 | 3.71E-02 | Sphingobacteriales |
| 8.45 | -0.21 | 7.11E-02 | Xanthomonadales |
| 0.32 | -0.29 | 1.22E-01 | Solirubrobacterales |
| 0.22 | 2.11 | 1.34E-01 | Ktedonobacterales |
| 0.40 | 0.41 | 1.81E-01 | Rhodocyclales |
| 0.14 | 0.36 | 1.83E-01 | Gaiellales |
| 0.25 | 0.38 | 2.02E-01 | Gallionellales |
| 1.30 | 1.54 | 2.10E-01 | Enterobacteriales |
| 11.59 | -0.04 | 2.43E-01 | Burkholderiales |
| 0.01 | -1.36 | 2.78E-01 | Gammaproteobacteria incertae sedis |
| 0.33 | 0.32 | 3.04E-01 | Gp1 |
| 0.02 | -1.22 | 3.07E-01 | Oligoflexales |
| 3.48 | -0.22 | 3.62E-01 | Flavobacteriales |
| 0.09 | 0.44 | 3.73E-01 | Nitrosomonadales |
| 0.36 | 0.39 | 3.88E-01 | Chlamydiales |
| 0.82 | 0.15 | 3.89E-01 | Opitutales |
| 0.12 | 0.24 | 4.07E-01 | Holophagales |
| 1.19 | -0.12 | 4.54E-01 | Verrucomicrobiales |
| 0.03 | -0.78 | 4.86E-01 | Candidatus_Koribacter |
| 0.02 | 0.55 | 4.87E-01 | Methanosarcinales |
| 3.09 | -0.12 | 5.01E-01 | Alphaproteobacteria incertae sedis |
| 0.01 | 1.09 | 5.05E-01 | Rhodobacterales |
| 3.11 | -0.14 | 5.23E-01 | Actinomycetales |
| 0.68 | -0.17 | 5.24E-01 | Bdellovibrionales |
| 0.25 | 0.08 | 5.34E-01 | Armatimonadales |
| 0.00 | 0.89 | 5.37E-01 | Sneathiellales |
| 0.01 | -1.02 | 5.37E-01 | Nitrososphaeraceae |
| 1.84 | 0.12 | 5.58E-01 | Cytophagales |
| 0.11 | -0.40 | 5.59E-01 | Gemmatimonadales |
| 0.41 | -0.12 | 5.62E-01 | Gp3 |
| 0.23 | -0.12 | 5.90E-01 | Candidatus_Solibacter |
| 0.01 | 0.64 | 5.94E-01 | Chthonomonadales |
| 1.55 | -0.07 | 6.14E-01 | Rhodospirillales |
| 0.02 | -0.61 | 6.15E-01 | Tepidisphaerales |
| 0.07 | 0.50 | 6.22E-01 | Neisseriales |
| 0.00 | -0.98 | 6.69E-01 | Methylococcales |
| 0.06 | 0.28 | 6.74E-01 | Bacillales |
| 1.42 | 0.08 | 6.79E-01 | Planctomycetales |
| 0.03 | 0.36 | 6.87E-01 | Terrimicrobium |
| 0.02 | 0.36 | 7.10E-01 | Gp2 |
| 0.31 | -0.36 | 7.12E-01 | Methylophilales |
| 0.00 | 0.82 | 7.14E-01 | Gp6 |
| 0.88 | -0.12 | 7.27E-01 | Chromatiales |
| 0.03 | 0.47 | 7.39E-01 | Acidimicrobiales |
| 0.35 | -0.10 | 7.69E-01 | Spirochaetales |
| 0.86 | 0.05 | 8.15E-01 | Telmatobacter |
| 3.71 | -0.02 | 8.76E-01 | Caulobacterales |
| 0.01 | -0.13 | 8.97E-01 | Lactobacillales |
| 0.01 | 0.13 | 9.13E-01 | Oceanospirillales |
| 1.26 | 0.02 | 9.29E-01 | Myxococcales |
| 0.00 | -0.14 | 9.37E-01 | Rickettsiales |
| 0.95 | -0.02 | 9.47E-01 | Terriglobus |
| 0.00 | 0.08 | 9.68E-01 | Gp14 |
| 0.02 | -0.01 | 9.88E-01 | Edaphobacter |
| 0.15 | -0.01 | 9.93E-01 | Acidobacterium |
| 9.35 | 0.00 | 9.93E-01 | Rhizobiales |

| ***V. dahliae* gene ID** | **Predicted structural homolog** | **Confidence (%)** | **Coverage (%)** | **Size (kDa)** | **Cysteines (%)** | **Isoelectric point** |
| --- | --- | --- | --- | --- | --- | --- |
| *VDAG_JR2_Chr2g00770* | Aerolysin/toxin | 82.6 | 55 | 23.7 | 4.6 | 5.74 |
| *VDAG_JR2_Chr3g05620* | Defensin | 95.6 | 81 | 7 | 9.5 | 6.98 |
| *VDAG_JR2_Chr3g12130* | Microbial ribonuclease | 100 | 84 | 10.8 | 3.3 | 8.57 |
| *VDAG_JR2_Chr4g10160* | Aerolysin/toxin | 92.7 | 22 | 42 | 2.1 | 4.77 |
| *VDAG_JR2_Chr5g07180* | Defensin | 59.9 | 21 | 9.4 | 4.5 | 8.87 |
| *VDAG_JR2_Chr5g09210* | Heparin-binding protein | 58.6 | 37 | 21.8 | 2.8 | 4.06 |
| *VDAG_JR2_Chr5g09660* | Aerolysin/toxin | 79.4 | 59 | 25.2 | 1.7 | 5.71 |
| *VDAG_JR2_Chr6g00350* | Leukocidin | 79.3 | 20 | 38.5 | 2.9 | 4.43 |
| *VDAG_JR2_Chr6g03080* | Defensin | 28.1 | 21 | 10.7 | 6.2 | 5.15 |
| *VDAG_JR2_Chr8g09620* | Defensin | 75.8 | 20 | 12.2 | 7.1 | 8.99 |

**Extended data Table 4: Effectors of *V. dahliae* strain JR2 with predicted structural homology with known antimicrobial proteins**

**Extended data Table 5: Sample statistics 16S ribosomal DNA profiling**

| **Sample name** | **Host** | **Tissue** | ***V. dahliae* infection** | **Quality filtered paired-end reads** | **Percentage of reads assigned to microbiome** |
| --- | --- | --- | --- | --- | --- |
| Sl_Mock_A | Tomato | Roots + rhizosphere | Mock | 41164 | 40.8 |
| Sl_Mock_B | Tomato | Roots + rhizosphere | Mock | 48069 | 46.4 |
| Sl_Mock_C | Tomato | Roots + rhizosphere | Mock | 43366 | 40.9 |
| Sl_WT_A | Tomato | Roots + rhizosphere | *V. dahliae* WT | 33386 | 41.2 |
| Sl_WT_B | Tomato | Roots + rhizosphere | *V. dahliae* WT | 39518 | 40.1 |
| Sl_WT_C | Tomato | Roots + rhizosphere | *V. dahliae* WT | 43502 | 41.6 |
| Sl_ΔVdAve1_A | Tomato | Roots + rhizosphere | *V. dahliae* *ΔVdAve1* | 51130 | 49.2 |
| Sl_ΔVdAve1_B | Tomato | Roots + rhizosphere | *V. dahliae ΔVdAve1* | 43869 | 43.5 |
| Sl_ΔVdAve1_C | Tomato | Roots + rhizosphere | *V. dahliae ΔVdAve1* | 46159 | 41.3 |
| Gh_Mock_A | Cotton | Roots + rhizosphere | Mock | 93241 | 48.2 |
| Gh_Mock_B | Cotton | Roots + rhizosphere | Mock | 90366 | 46.1 |
| Gh_Mock_C | Cotton | Roots + rhizosphere | Mock | 92144 | 43.8 |
| Gh_WT_A | Cotton | Roots + rhizosphere | *V. dahliae* WT | 115812 | 49.6 |
| Gh_WT_B | Cotton | Roots + rhizosphere | *V. dahliae* WT | 105032 | 53.5 |
| Gh_WT_C | Cotton | Roots + rhizosphere | *V. dahliae* WT | 91601 | 45.9 |
| Gh_ΔVdAve1_A | Cotton | Roots + rhizosphere | *V. dahliae* *ΔVdAve1* | 92975 | 48.1 |
| Gh_ΔVdAve1_B | Cotton | Roots + rhizosphere | *V. dahliae* *ΔVdAve1* | 118244 | 51.6 |
| Gh_ΔVdAve1_C | Cotton | Roots + rhizosphere | *V. dahliae* *ΔVdAve1* | 112762 | 50.4 |
| Sl_Sm_WT_A | Tomato | Stems | *V. dahliae* WT | 85526 | 3.0 |
| Sl_Sm_WT_B | Tomato | Stems | *V. dahliae* WT | 87981 | 2.5 |
| Sl_Sm_WT_C | Tomato | Stems | *V. dahliae* WT | 71022 | 0.6 |
| Sl_Sm_ΔVdAve1_A | Tomato | Stems | *V. dahliae* *ΔVdAve1* | 110437 | 3.2 |
| Sl_Sm_ΔVdAve1_B | Tomato | Stems | *V. dahliae* *ΔVdAve1* | 112487 | 1.9 |
| Sl_Sm_ΔVdAve1_C | Tomato | Stems | *V. dahliae* *ΔVdAve1* | 115161 | 1.5 |

**Extended data Table 6: Primers used in this study**

| **Name** | **Sequence (5' --> 3')** | **Application** |
| --- | --- | --- |
| XhoI_VdAve1_Fw | CGGTATCTCGAGGATCTAGGGACCGCATCCTAC | Protein production |
| VdAve1_BamHI_Rv | CGTCTAGGATCCTCATTATTATATCTGTCTAAATTCGATGTTGACC | Protein production |
| XhoI_VnAve1_Fw | CGGTATCTCGAGCAATTAGGGACCGCATCC | Protein production |
| VnAve1_BamHI_Rv | CGTCTAGGATCCTCATTATTATATCTGTTCAAACTCG | Protein production |
| VdAve1_qPCR_Fw | TGTTACCAAAGCAGCACACAAGG | Real-time PCR |
| VdAve1_qPCR_Rv | CCTTATGCCTCGTTCCCTTCCAC | Real-time PCR |
| VdGAPDH_Fw | CGAGTCCACTGGTGTCTTCA | Real-time PCR |
| VdGAPDH_Rv | CCCTCAACGATGGTGAACTT | Real-time PCR |
| ITS1-Fw | AAAGTTTTAATGGTTCGCTAAGA | Real-time PCR |
| St-Ve1-Rv | CTTGGTCATTTAGAGGAAGTAA | Real-time PCR |
| SlRUB_Fw | GAACAGTTTCTCACTGTTGAC | Real-time PCR |
| SlRUB_Rv | CGTGAGAACCATAAGTCACC | Real-time PCR |
| AtRUB_Fw | GCAAGTGTTGGGTTCAAAGCTGGTG | Real-time PCR |
| AtRUB_Rv | CCAGGTTGAGGAGTTACTCGGAATGCTG | Real-time PCR |
| Novosphingobium_Fw | CGGAATAACAGTTAGAAATGACTGC | Real-time PCR |
| Novosphingobium_Rv | CAGGTACTGTCATTATCATCCCT | Real-time PCR |
| Sphingopyxis_Fw | CGGAATAACTCAGAGAAATTTGTGC | Real-time PCR |
| Sphingopyxis_Rv | CCGGTACTGTCATTATCATCCCG | Real-time PCR |
| VdAMP2_qPCR_Fw | CATGGAACACGACCAACTGC | Real-time PCR |
| VdAMP2_qPCR_Rv | GATCCATTCGGGCTTCGACT | Real-time PCR |
| 16s_Bact340F | TCCTACGGGAGGCAGCAGT | Real-time PCR |
| 16s_533R | TTACCGCGGCTGCTGGCAC | Real-time PCR |
| PacI_VdAMP2_FT_Fw | CGGTATTTAATTAAATGGCCACCCGCCGAAC | To generate pVdAve1::VdAMP2 transformant |
| VdAMP2_NotI_RV | CGTCTAGCGGCCGCTCAGATCGAGAAGCCGAG | To generate pVdAve1::VdAMP2 transformant |
| JR2_VdAMP2_LB_Fw | GGTCTTAAUAGGCGTAGGGAGGAGATGTT | To generate VdAMP2 deletion mutant |
| JR2_VdAMP2_LB_Rv | GGCATTAAUCCATGCTAGGCACAGACCAA | To generate VdAMP2 deletion mutant |
| JR2_VdAMP2_RB_Fw | GGACTTAAUATTCGTGTGAAGCCCTTGGA | To generate VdAMP2 deletion mutant |
| JR2_VdAMP2_RB_Rv | GGGTTTAAUGTTGCAGAAGCCCTGTTTGG | To generate VdAMP2 deletion mutant |
| 27F | AGAGTTTGATCCTGGCTCAG | 16S identification |
| 1492R | GGWTACCTTGTTACGACT | 16S identification |
